# Kill and cure: genomic phylogeny and bioactivity of a diverse collection of *Burkholderia gladioli* bacteria capable of pathogenic and beneficial lifestyles

**DOI:** 10.1101/2020.04.09.033878

**Authors:** Cerith Jones, Gordon Webster, Alex J. Mullins, Matthew Jenner, Matthew J. Bull, Yousef Dashti, Theodore Spilker, Julian Parkhill, Thomas R. Connor, John J. LiPuma, Gregory L. Challis, Eshwar Mahenthiralingam

**Affiliations:** Microbiomes, Microbes and Informatics Group, Organisms and Environment Division, School of Biosciences, Cardiff University, Cardiff, Wales, CF10 3AX, UK; Department of Chemistry, University of Warwick, CV4 7AL, UK; Department of Pediatrics, University of Michigan Medical School, Ann Arbor, Michigan, USA; Wellcome Trust Sanger Institute, Wellcome Trust Genome Campus, Hinxton, Cambridge, CB10 1SA UK; School of Applied Sciences, Faculty of Computing, Engineering and Science, University of South Wales, Pontypridd, CF37 4BD, UK; Pathogen Genomics Unit, Public Health Wales Microbiology Cardiff, University Hospital of Wales, Cardiff, CF14 4XW; The Centre for Bacterial Cell Biology, Biosciences Institute, Medical School, Newcastle University, Newcastle upon Tyne, NE2 4AX, UK; Department of Veterinary Medicine, Madingley Road, Cambridge, CB3 0ES; Warwick Integrative Synthetic Biology Centre, University of Warwick, Coventry CV4 7AL, UK; Department of Biochemistry and Molecular Biology, Biomedicine Discovery Institute, Monash University, Clayton, VIC 3800, Australia

**Author notes:** Corresponding Author: Prof. Eshwar Mahenthiralingam, Microbiomes, Microbes and Informatics Group, Organisms and Environment Division, School of Biosciences, Cardiff University, Sir Martin Evans Building, Room 2.06, Museum Avenue, Cardiff CF10 3AX, United Kingdom. Co-corresponding Author: Cerith Jones, School of Applied Sciences, Faculty of Computing, Engineering and Science, University of South Wales, Pontypridd, CF37 4BD, United Kingdom.

**Keywords:** *Burkholderia*, *B. gladioli*, antibiotic production, plant pathogenesis, cystic fibrosis infection, phylogenomics

## Abstract

*Burkholderia gladioli* is one of few bacteria with a broad ecology spanning disease in humans, animals, and plants, and encompassing beneficial interactions with multiple eukaryotic hosts. It is a plant pathogen, a bongkrekic acid toxin producing food-poisoning agent, and a lung pathogen in people with cystic fibrosis (CF). Contrasting beneficial traits include antifungal production exploited by insects to protect their eggs, plant protective abilities and antibiotic biosynthesis. We explored the ecological diversity and specialized metabolite biosynthesis of 206 *B. gladioli* strains, phylogenomically defining 5 evolutionary clades. Historical disease pathovars (pv) *B. gladioli* pv. *allicola* and *B. gladioli* pv. *cocovenenans* were phylogenetically distinct, while *B. gladioli* pv. *gladioli* and *B. gladioli* pv. *agaricicola* were indistinguishable. Soft-rot disease and CF infection pathogenicity traits were conserved across all pathovars. Biosynthetic gene clusters for toxoflavin, caryoynencin and enacyloxin were dispersed across *B. gladioli*, but bongkrekic acid and gladiolin production were clade specific. Strikingly, 13% of CF-infection strains characterised (n=194) were bongkrekic acid toxin positive, uniquely linking this food-poisoning risk factor to chronic lung disease. Toxin production was suppressed by exposing strains to the antibiotic trimethoprim, providing a potential therapeutic strategy to minimise poisoning risk in CF.

## INTRODUCTION

The genus *Burkholderia* contains important plant, animal and human pathogenic bacteria (Depoorter et al 2016, Lipuma 2010) as well as environmentally beneficial species (Suarez-Moreno et al 2012). Recently, amino acid and nucleotide based analyses have split *Burkholderia* strains into five distinct lineages corresponding to *Burkholderia* sensu stricto, *Paraburkholderia, Caballeronia, Robbsia, Trinickia* and *Mycetohabitans*, and a single species lineage represented by *Paraburkholderia rhizoxinica* (Estrada-de Los Santos et al 2018). Within *Burkholderia* sensu stricto, the *Burkholderia cepacia* complex group of species are problematic lung pathogens in people with cystic fibrosis (CF) (Lipuma 2010). The three most commonly isolated *Burkholderia* species among US CF patients are *B. multivorans, B. cenocepacia*, and interestingly, *B. gladioli* (Lipuma 2010). Although phenotypically similar, genetically *B. gladioli* is not a member of the *B. cepacia* complex, but is part of a group of species associated with plant disease including *B. glumae* and *B. plantarii* (Suarez-Moreno et al 2012). In relation to CF infection, *B. gladioli* may cause severe systemic abscesses (Jones et al 2001) and is also considered a risk factor for lung transplantation since it is associated with poor clinical outcome (Murray et al 2008). While the potential for patient-to-patient spread and rapid clinical decline are identified traits of *B. cepacia* complex infection in people with CF (Lipuma 2010), the population biology, epidemiology and genomics of *B. gladioli* as a lung pathogen are essentially unknown.

In relation to its environmental ecology, *B. gladioli* was originally isolated as a pathogen of the *Gladiolus* genus of flowering plants and its taxonomy updated several times (Coenye et al 1999). The current species encompasses the historical *Gladiolus*-disease causing taxa “*Pseudomonas gladioli”* and *“Pseudomonas marginata”* (Hildebrand et al 1973), the food poisoning-associated *B. cocovenenans* (Coenye et al 1999), and the potential biological control agent “*Pseudomonas antimicrobica”* (Coenye et al 2000). *B. gladioli* has also been isolated as a pathogen of important crops that resulted in pathovar (pv.) designations being applied to the causative isolates of: mushroom rot, *B. gladioli* pv. *agaricicola* (Gill and Tsuneda 1997); onion rot, *B. gladioli* pv. *allicola* (Wright et al 1993); and the historical bulb rot disease, *B. gladioli* pv. *gladioli* (Hildebrand et al 1973). *B. gladioli* and its close relative *B. glumae* are also major rice pathogens causing panicle blight (Nandakumar et al 2009). *B. cocovenenans* represents a fourth pathovar (Jiao et al 2003) that is responsible for food poisoning when tempe bongkrek, the fermented coconut-based Indonesian national dish, is produced with *Rhizopus* fungal cultures contaminated with *B. gladioli* (Moebius et al 2012). Under these conditions, a polyketide biosynthetic gene cluster (BGC) is activated resulting in *B. gladioli* pv. *cocovenenans* producing the respiratory toxin bongkrekic acid that is fatal when ingested (Moebius et al 2012). The *B. gladioli* pathovars had been assigned based on the source of isolates and researchers have argued that there is a need to differentiate the lethal toxin producing pathovars such as *B. gladioli* pv. *cocovenenans* (Jiao et al 2003). However, the evolutionary basis of the pathovar designations of *B. gladioli* remains to be systematically investigated.

The capacity to produce a diverse range of specialized metabolites ranging from toxins such as bongkrekic acid (Moebius et al 2012) to beneficial antibiotics is a common trait among *Burkholderia* bacteria (Depoorter et al 2016, Kunakom and Eustaquio 2019). Close ecological associations with multiple eukaryotic hosts is a key primer for metabolite production by *B. gladioli*. As a detrimental trait, it produces the bright yellow phytotoxin, toxoflavin, which enhances the virulence of *B. gladioli* in rice disease (Lee et al 2016). In parallel with bongkrekic acid biosynthesis, *B. gladioli* also produces the polyketide enacyloxin in co-culture with the fungus *Rhizopus microspores* (Ross et al 2014a). A close association of *B. gladioli* with fungi was linked to the discovery that the bacterium encodes a nonribosomal peptide synthetase BGC that produces icosalide A1, a metabolite originally characterised as a product of an *Aureobasidium* fungus (Dose et al 2018, Jenner et al 2019). PCR screening of DNA extracts from the original *Aureobasidium* culture demonstrated that a *B. gladioli* co-culture was present (Jenner et al 2019). The vertical transmission of symbiotic *B. gladioli* in herbivorous *Lagriinae* beetles clearly demonstrates how ecological benefit may derive from the metabolites the bacterium produces (Florez et al 2017). *B. gladioli* was found in the reproductive tract of the beetles and resulted in the *Lagriinae* eggs gaining protection from fungal attack by the antimicrobial bacterial metabolites including toxoflavin, caryoynencin, the novel polyketide lagriene and novel aromatic glycoside sinapigladioside (Florez et al 2017).

Genomics has revolutionized our understanding of *Burkholderia* population biology, and the beneficial and detrimental interactions of these ecologically diverse bacteria. The discovery of gladiolin, a novel polyketide antibiotic with promising activity against *Mycobacterium tuberculosis*, was greatly enhanced by complete genome sequencing of strain *B. gladioli* BCC0238 (Song et al 2017). Using a combination of systematic approaches including genome mining for specialized metabolite BGCs, metabolite characterisation and phenotypic assays, production of the antimicrobial cepacin was shown to underpin biological control of damping-off disease by the biopesticide species *Burkholderia ambifaria* (Mullins et al 2019). However, a limited number of complete genome sequences are available for *B. gladioli* which include those for strain BSR3, a rice disease isolate (Seo et al 2011), the bulb-associated Type strain ATCC 10248 (Johnson et al 2015), and the CF lung infection isolate, BCC0238 (Song et al 2017). Here we investigate the population biology of *B. gladioli* as a functionally diverse species that interacts with human, plant, insect and microbial ecosystems. Using genome sequence analysis of 206 *B. gladioli* strains from diverse sources we defined the genetic linkage to pathovar status, mapped the ability to mediate plant soft-rot and human disease, and correlated population biology to capacity for specialized metabolite production. The genomics-based taxonomy of all the *B. gladioli* isolates was consistent with their designation as a single species. Pathovars *B. gladioli* pv. *allicola* and *B. gladioli* pv. *cocovenenans*, as well as biosynthetic clusters for bongkrekic acid and gladiolin, were shown to be clade restricted within the overall *B. gladioli* population. People with CF were susceptible to all clades of *B. gladioli* and the presence of the bongkrekic acid BGC was revealed as a new risk factor for these infectious isolates.

## MATERIALS AND METHODS

### Bacterial strains and growth conditions

A collection of 206 *B. gladioli* isolates was assembled for this study and their source details, genomic features and the analysis they were subject to are included in supplementary data (Table S1). These were drawn from the Cardiff collection (Mahenthiralingam et al 2011, Mullins et al 2019, Song et al 2017) and *Burkholderia cepacia* Research Laboratory and Repository (University of Michigan, Michigan, USA) (Lipuma 2010), with additional reference and pathovar strains of *B. gladioli* obtained from the Belgium Coordinated Collection of Microorganisms (Ghent, Belgium) and National Collections of Plant Pathogenic Bacteria (York, United Kingdom) (Table S1). *B. gladioli* isolates were routinely grown on Tryptone Soya Agar (TSA) or within Tryptone Soya Broth (TSB) liquid cultures, and incubated at 37°C. Antibiotic production was induced by growing strains on a minimal salts media with glycerol as the sole carbon source (designated Basal Salts Medium with Glycerol; BSM-G) as previously described (Hareland et al 1975, Mahenthiralingam et al 2011).

Antimicrobial antagonism assays were performed by overlaying with the following susceptibility testing organisms: *Staphylococcus aureus* ATCC25923, *Ralstonia mannitolilytica* LMG6866, and *Candida albicans* SC 5314 as previously described (Mahenthiralingam et al 2011). *Escherichia coli* strain NCTC 12241 was used as a control for the mushroom and onion rot assays. Artificial CF sputum medium was made up as previously described (Kirchner et al 2012) to model if bongkrekic acid production occurred under CF lung infection-like growth conditions. Trimethoprim (1 µg/mL) was incorporated into BSM-G to determine if *B. gladioli* metabolite production was induced by sub-inhibitory concentrations of this antibiotic as described for *B. thailandensis* (Okada et al 2016).

### Genome sequencing, assembly and analysis

Genomic DNA was prepared from 3 mL TSB overnight cultures of *B. gladioli*. Cells were harvested by centrifugation and suspended in 400 µl of 4 M guanidine isothiocyanate solution (Invitrogen, UK). DNA was extracted from these bacterial suspensions using a Maxwell^®^ 16 automated nucleic acid purification system and the Maxwell^®^ tissue DNA purification kit following the manufacturer’s instructions (Promega, UK). Purified DNA extracts were treated with RNase (New England BioLabs, UK). Genomes were sequenced using the Illumina HiSeq 2000 and HiSeq X Ten platforms at the Wellcome Sanger Institute as previously described (Mullins et al 2019). Genomes were assembled from the read data, annotated and compared using a virtual machine hosted by the Cloud Infrastructure for Microbial Bioinformatics (CLIMB) consortium (Connor et al 2016). Sequence reads were trimmed using Trim Galore v0.4.2 (Babraham Bioinformatics), overlapped using FLASH v1.2.11 (Magoc and Salzberg 2011), and assembled using SPAdes v 3.9.1 (Bankevich et al 2012). Assembled genomes were polished using Pilon v1.21 (Walker et al 2014). Prokka v 1.12 beta (Seemann 2014) was used for gene prediction and annotation. The quality of genome assemblies was assessed using Quast (Gurevich et al 2013) and Prokka annotations cross-compared with gene predictions generated by Glimmer v3.02b (Delcher et al 2007). Draft genome contigs were ordered against a complete reference genome for *B. gladioli* BCC0238 (Song et al 2017) using CONTUGuator v2.7 (Galardini et al 2011). To supplement the Illumina sequencing, contiguated genomes were generated for clade specific strains BCC1710, BCC1621 and BCC1622 (see Table S1) using Pacific Biosciences Single Molecule Real Time sequencing as described (Song et al 2017).

Average nucleotide identity (ANI) was used for genomic taxonomy and calculated using PyANI v0.2.1 (Pritchard et al 2016). The *B. gladioli* core genome was computed using Roary v3.6.0 (Page et al 2015). Maximum-likelihood trees were drawn from the core gene alignment with FastTree (Price et al 2010) using the generalised time-reversible model of nucleotide evolution and visualised using FigTree (http://tree.bio.ed.ac.uk/software/figtree). Rooting the trees with multiple *Burkholderiales* species (*Burkholderia glumae, Burkholderia oklahomiensis* and *Paraburkholderia xenovorans*) failed to produce a biologically meaningful root. The closest sequences to these outgroups was variable but all produced trees of consistent phylogenetic separation for the *B. gladioli* clades identified. Therefore, an unrooted tree was presented in the final analysis. *B. gladioli* MLST sequence type assignments were made by using the PubMLST database and website (Jolley and Maiden 2010), via the MLST tool developed by Torsten Seemann (https://github.com/tseemann/mlst). Initial profiling of specialized metabolite biosynthetic gene potential was predicted for *B. gladioli* genomes via antiSMASH (Blin et al 2017, Weber et al 2015) running as a local instance on CLIMB. The presence or absence of known BGCs was determined by mapping sequencing reads to representative BCG reference sequences using snippy (https://github.com/tseemann/snippy). Percentage of reads mapping to the reference sequence, and the actual number of corresponding reads were used to manually determine the status of each BGC in a given strain.

### Mushroom and onion rot bioassays

Mushroom (*Agaricus bisporus*) soft-rot bioassays were carried out as described (Roy Chowdhury and Heinemann 2006) with the surface sterilisation and immersion into ice-cold water step omitted as this caused non-specific rotting of mushrooms. Briefly, mushrooms (Oakland closed cup mushroom, Lidl UK GmbH, produced in Ireland) were cut into 3-4 mm slices with a sterile blade. *B. gladioli* was grown overnight in TSB and the cap of each mushroom was inoculated with a 10 µl drop of bacterial suspension adjusted to 0.1 OD600 nm in TSB. Onion (*Allium cepa*) soft-rot bioassays were carried out as described (Jacobs et al 2008). Brown onions (Tesco, Cardiff, UK) had their skin and outer onion layer removed prior to quartering with a sterile knife. Individual onion layers were cut into 3 to 4 cm pieces, wounded on their inner surface with a knife slit made under aseptic conditions, and the wound inoculated with 10 µl of bacterial suspension made as described for the mushroom assay. All assays (the test *B. gladioli*, a control *E. coli* NCTC 12241 strain, a TSB control and untreated controls) were performed in triplicate on sterile wet filter paper contained in sterile 9 cm plastic Petri dishes, sealed with Parafilm M and incubated at 30°C for 48 h.

### Preparation of *B. gladioli* metabolite extracts and antimicrobial activity

To analyse the metabolites produced by different *B. gladioli* strains, BSM-G agar plates (5 per strain) were streaked with cells from a freshly revived culture and incubated for 72 h at 30°C. Bacterial growth was removed using a sterile cell scraper and the spent agar was transferred to a glass bottle. Metabolites were extracted from the agar using dichloromethane (2 h with gentle shaking). The crude extract was concentrated to dryness under a vacuum at 22°C and resuspended in 1 mL of dichloromethane. The bioactivity of each extract and control dichloromethane was tested by pipetting 5 µL onto a TSA plate and allowing the plates to dry and solvent to evaporate. Each plate was then overlaid with molten Iso-sensitest agar (Oxoid, UK) seeded with *S. aureus, R. mannitolilytica* or *C. albicans* as described (Mahenthiralingam et al 2011). Plates were incubated at 37°C for 24 h and photographed to document the inhibitory zones of clearing observed. Bioactivity assays were performed in triplicate for each strain.

### Analysis of *B. gladioli* metabolites by high performance liquid chromatography (HPLC)

HPLC analysis was used to quantify and identify known *B. gladioli* metabolites as follows. Specialized metabolites were induced using growth on BSM-G media (as above) and extracted directly from a 20 mm agar disc cut from the plates as described (Mullins et al 2019). Extracts (20 µL injection volume) were analysed on a Waters® AutoPurification™ HPLC System fitted with a reverse phase analytical column (Waters® XSelect CSH C18, 4.6 x 100 mm, 5 μm) and a C18 SecurityGuard™ cartridge (Phenomenex) in series. Absorbance at 210-400 nm was monitored by a photo diode array detector (PDA). Mobile phases consisted of A: water with 0.1% formic acid and B: acetonitrile with 0.1% formic acid. A flow rate of 1.5 ml min^−1^ was used. Elution conditions were as follows. 0-1 minutes: 95% A / 5% B; 1-9 minutes: gradient of A from 95 to 5% / gradient of B from 5% to 95%; 10-11 minutes: 5% A / 95% B; 11-15 minutes: 95% A / 5% B. Peak height and area were calculated using MassLynx V4.1 software (www.waters.com).

To enable further biosynthetic pathway-metabolite correlations, *B. gladioli* gene mutants were used for the gladiolin BGC (Song et al 2017), and constructed *de novo* for the toxoflavin, bongkrekic acid and caryoynencin pathways as follows. PCR products encoding fragments of core biosynthetic genes were amplified using specific primers (see Table S2) and cloned into the pGpΩTp suicide plasmid (Flannagan et al 2007) following digestion with *Xba*I/*Eco*RI (*bonA* and *cayA*), or *Xba*I/*Kpn*I (*toxA*). Plasmids were mobilised as described (Song et al 2017) and mutants created in *B. gladioli* BCC0238 for the gladiolin and toxoflavin BGCs, strain BCC1710 for the bongkrekic acid BGC and strain BCC1697 for the caryoynencin pathway. Comparative analysis of metabolite extracts from parental versus mutant strains for each of the latter BGCs, together with correlation to mass spectrometry analysis (see below), was used to identify HPLC metabolite peaks.

### Culture conditions, extraction protocol and high-resolution mass spectrometry

Known *Burkholderia* metabolites were confirmed by mass spectrometry essentially as described (Jenner et al 2019, Mahenthiralingam et al 2011, Mullins et al 2019, Song et al 2017). Briefly, all *B. gladioli* strains were grown at 30 °C on BSM-G plates. Single plates were extracted by removal of the cell material, chopping of the agar and addition of 4 mL ethyl acetate for 2 h. Centrifugation in a 1.5 mL Eppendorf tube was used to remove debris. Crude extracts were directly analysed by UHPLC-ESI-Q-TOF-MS. UHPLC-ESI-Q-TOF-MS analyses were performed using a Dionex UltiMate 3000 UHPLC connected to a Zorbax Eclipse Plus C-18 column (100 × 2.1 mm, 1.8 μm) coupled to a Bruker MaXis II mass spectrometer. Mobile phases consisted of water (A) and acetonitrile (B), each supplemented with 0.1% formic acid. A gradient of 5 % B to 100 % B over 30 min was used at a flow rate of 0.2 ml min^−1^. The mass spectrometer was operated in either positive- or negative-ion mode with a scan range of 50– 3,000 *m/z*. Source conditions were: end-plate offset at −500 V, capillary at −4,500 V, nebulizer gas (N2) at 1.6 bar, dry gas (N2) at 8 L min^−1^ and dry temperature at 180 °C. Ion transfer conditions were: ion funnel radio frequency (RF) at 200 Vpp, multiple RF at 200 Vpp, quadrupole low mass at 55 *m/z*, collision energy at 5.0 eV, collision RF at 600 Vpp, ion cooler RF at 50–350 Vpp, transfer time at 121 μs and pre-pulse storage time at 1 μs. Calibration was performed with 1 mM sodium formate through a loop injection of 20 μl at the start of each run.

### PCR detection of the bongkrekic Acid BGC

To detect the presence of the bongkrekic acid BGC, PCR probes were designed to target the central polyketide synthase enzyme gene, *bonA*, within the gene cluster of *B. gladioli* BCC1710 (*bonA*-F, 5’ ATTTCTAGAAGTATCCGCATTTTCGTCGC 3’; *bonA*-R 5’ TATGAATTCGATCGATCAGTTGCGCTTCC 3’). PCRs were performed using the Taq PCR Core Kit (Qiagen) as per the manufacturer’s instructions and incorporating Q-solution.

Thermal cycling conditions used an annealing temperature of 54.5°C and extension time of 1 min 5 sec, run over 30 cycles. The 1053 bp *bonA* gene amplicon product was detected by gel electrophoresis and subjected to Sanger sequencing (Eurofins, Genomics) to confirm its identity from the control strain BCC1710.

### Accession Numbers

The sequencing read data of *B. gladioli* isolates from this study are available from the European Nucleotide Archive under the project accession number PRJEB9765; isolate accession numbers are provided in Table S1.

## RESULTS

### Assembly and genomic taxonomy of a *B. gladioli* isolate collection

To provide a holistic understanding of taxonomy and pathovar population biology of *B. gladioli*, a representative collection of 206 isolates was assembled and their genomes sequenced (Table S1). The majority of isolates (*n*=194) were from people with CF, with 181 from the USA, 7 from the UK, 4 from Canada, and one each from Australia and Italy (Table S1). Twelve strains were from environmental sources including pathovar reference isolates as follows: isolates of plant-disease associated *B. gladioli* pv. *gladioli* (*n*=3), pv. *agaricicola* (*n*=3) and pv. *alliicola* (n=3), and *B. gladioli* pv. *cocovenenans* (*n*=2) reference toxin producing strains. One *B. gladioli* isolated from an environmental industrial source was also included (BCC1317; Table S1). Short read genome sequencing yielded high quality draft genomes (average of 82 contigs, ranging from 20 [BCC1721] to 284 [BCC1788]) with a mean size for *B. gladioli* of 8.28 Mb, GC content of 68% and encoding a mean of 6872 protein-encoding genes (Table 1). These metrics were consistent with previously reported *B. gladioli* genomes (Johnson et al 2015, Seo et al 2011, Song et al 2017).

**Table 1.** Summary genome sequencing statistics

|  | Mean | Maximum | Minimum |
| --- | --- | --- | --- |
| Number of Contigs | 82.6 | 284 (BCC1788) | 20 (BCC1721) |
| Genome Size | 8.28 Mb | 8.94 Mb (BCC1815) | 7.32 Mb (BCC1681) |
| N50 | 404821 | 1964774 (BCC1691) | 147146 (BCC1713) |
| %GC | 68.0 % | 68.31 % (BCC1823) | 67.39 % (BCC1815) |
| Number of genes | 6872 | 7693 | 6012 |

Since the assignment of isolates within the *Burkholderia* genus (Estrada-de Los Santos et al 2018) generally, and *B. gladioli* specifically (Coenye et al 1999, Coenye et al 2000) have undergone multiple rounds of taxonomic re-classification, we initially established if the 206 *B. gladioli* isolates in the collection comprised a single bacterial species. Using average nucleotide identity, the 96.85% ANI for the entire *B. gladioli* 206 genome dataset was above the 95% identity required for designation as a single species (Goris et al 2007). This confirmed that the previous incorporation of “*P. cocovenenans”* (strains LMG 11626 and LMG 18113) (Coenye et al 1999) and “*P. marginata”* (ATCC10248) (Hildebrand et al 1973) into *B. gladioli* are supported by the genomic taxonomy (Goris et al 2007) (Table S1).

In addition, ANI heatmap analysis also suggested that a significant subspecies population structure existed within *B. gladioli* (Figure 1), and as such the following designation of groups was made. Group 1 (*n*=27) comprised 3 closely related sub-groups: 1A, containing the reference *B. gladioli* pv. *cocovenenans* strains, 1B and 1C; each subgroup was distinct in terms of their ANI relatedness (Figure 1). All isolates within each subgroup encoded the bongkrekic acid BGC (see below; Table S1) setting them apart from the rest of the *B. gladioli* collection and supporting their collective designation as Group 1. Group 2 was composed of 73 strains and included all 3 *B. gladioli* pv. *allicola* reference isolates. Group 3 (*n* = 106) contained both the *B. gladioli* pv. *agaricicola* and *B. gladioli pv. gladioli* reference isolates (Figure 1). Within each of these 3 initial groupings, the genomic ANI ranged from >98.1% (Group 3) to >99.1% (Group 1B), which was greater than the 96.85% collection average and suggested that distinct genetic lineages were present within *B. gladioli* (Figure 1).

**Figure 1.**
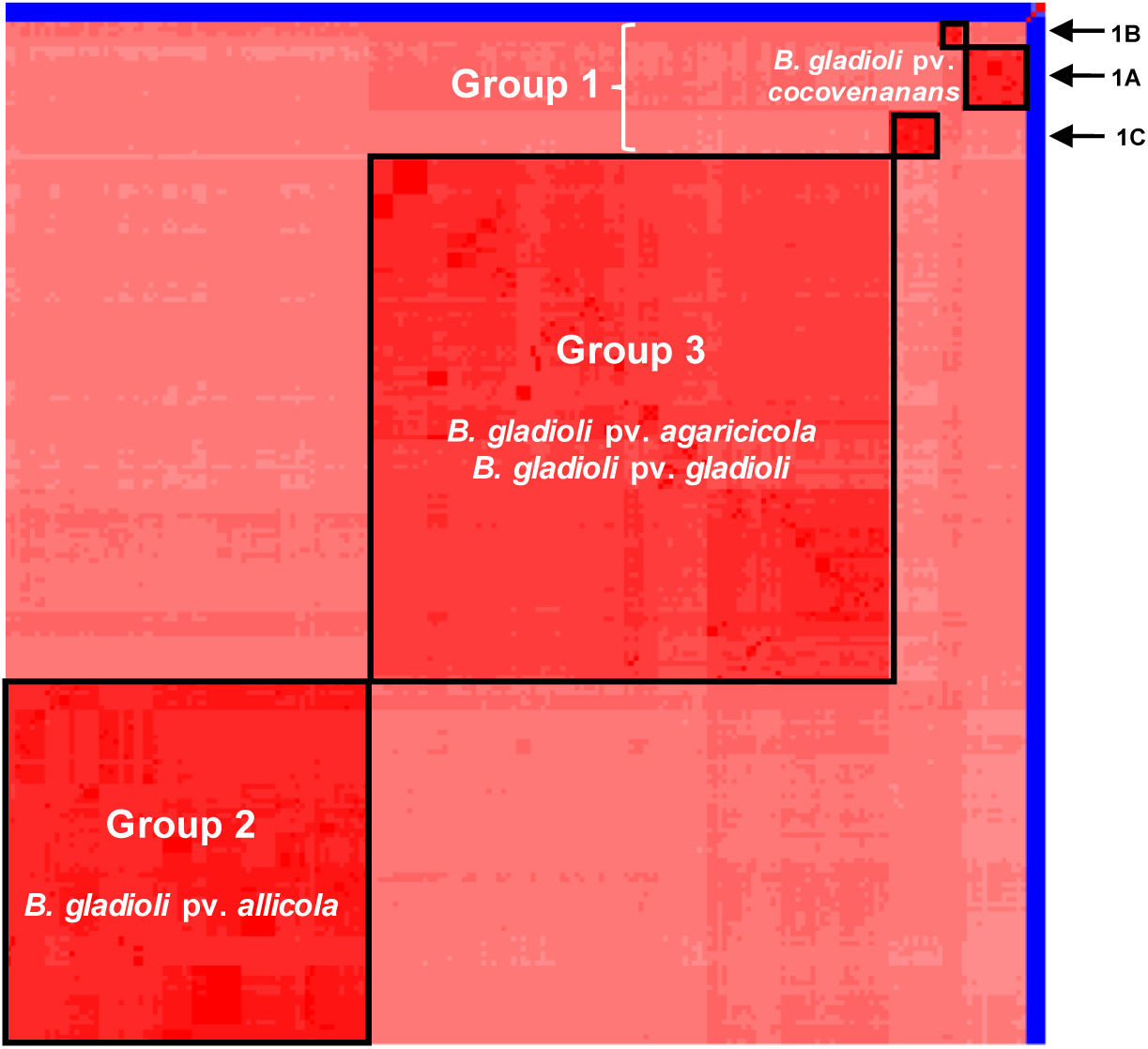
*B. gladioli* comprises a single genomic species with evidence of subspecies clustering by average nucleotide identity. The ANI of 206 *B. gladioli* genomes was compared using pyANI and a heatmap constructed (see methods). Darker red shading correlates to the greater percentage identity of each isolate. Subgroups with greater than 98.8% ANI are shown within the black outlined boxes. The main clusters are labelled as Group 1, 2 and 3, with Group 1 isolates sub-dividing further into sub-groups 1A, 1B and 1C (see top right). Four non-*B. gladioli* species genomes were used as taxonomic controls: 3 strains of the closely related species *B. glumae* and 1 strain of *B. ambifaria;* their low nucleotide identity to *B. gladioli* (ANI < 95%) is shown in blue.

### Core gene phylogenomic analysis reveals distinct evolutionary clades within *B. gladioli*

To investigate the evolutionary linkages behind the ANI groupings (Figure 1), we constructed a phylogeny from the 4406 core genes identified within the 206 *B. gladioli* genome dataset. The strain groups defined by ANI (Figure 1) were also supported as distinct evolutionary clades in the phylogenomic analysis (Figure 2). The three Group 1 ANI sub-clusters correspondingly separated as clades 1A, 1B and 1C, with the reference *B. gladioli* pv. *cocovenenans* strains locating specifically to clade 1A (Figure 2). These Group 1 strains separated as 13 isolates in clade 1A, 4 in clade 1B and 10 isolates in clade 1C (Table S1). At the distal ends of the *B. gladioli* phylogenetic tree were clade 2 and clade 3 strains (Figure 2), that also corresponded exactly with the respective ANI groupings (Figure 1). All three reference *B. gladioli* pv. *allicola* strains mapped to clade 2 suggesting that this pathovar status was evolutionarily supported, but *B. gladioli* pv. *agaricicola* and pv. *gladioli* grouped within clade 3 and were not genetically distinguished (except that they were distinct from clades 1 and 2; Figure 2).

**Figure 2.**
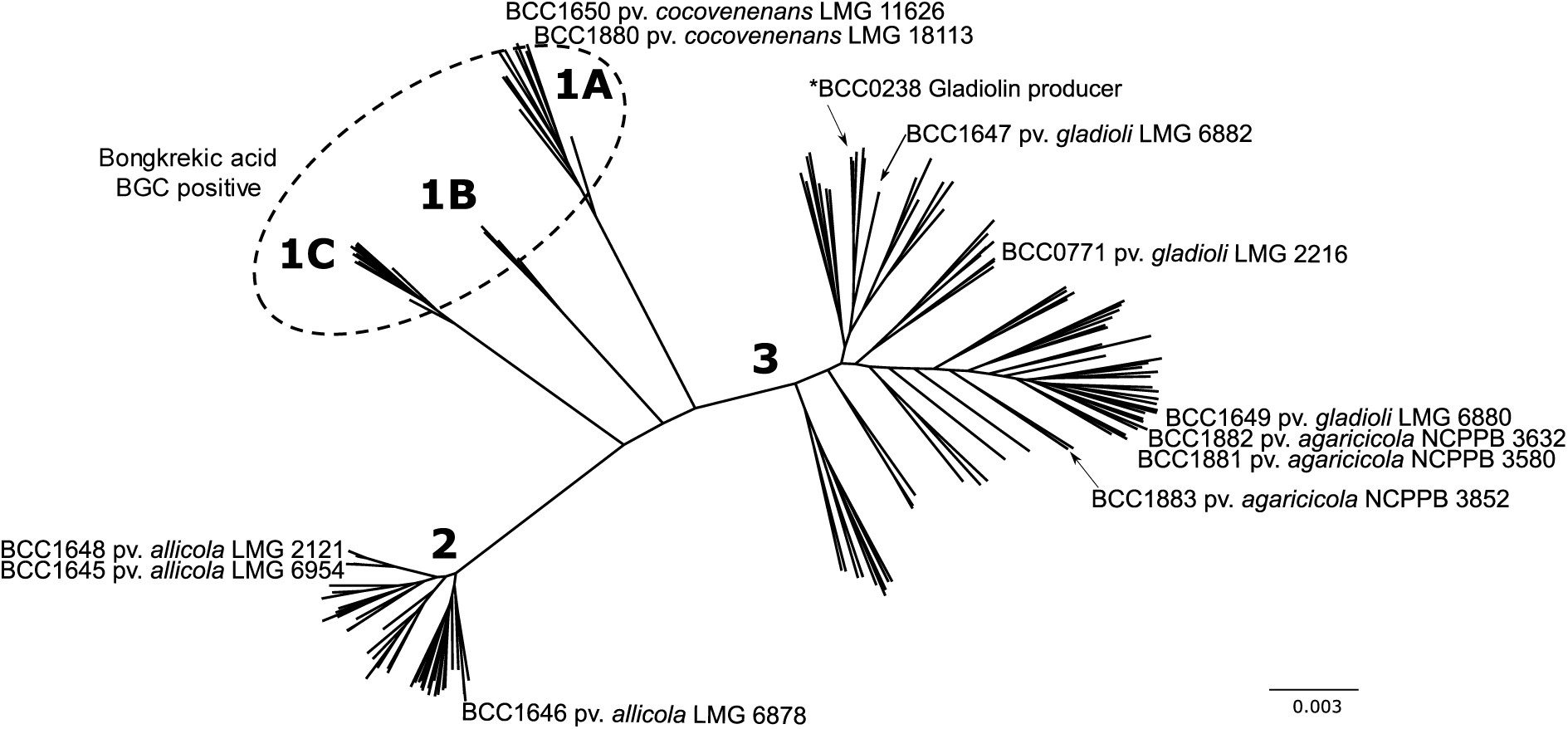
The population biology of *B. gladioli* inferred by core genome gene phylogeny. The core genome of the 206 *B. gladioli* strains were analysed using Roary and the resulting 4406 core genes aligned and used to construct a phylogeny. A maximum-likehood un-rooted tree was constructed using the generalised time-reversible model of nucleotide evolution; multiple rooting with species outside of *B. gladioli* was used to confirmed topology (see materials and methods). The position of the *B. gladioli* plant pathovar reference strains and model gladiolin-producing strain, BCC0238 is shown. The 5 major evolutionary branches consistent with the ANI groups and sub-groups are numbered accordingly. Clades 1A,1B and 1C all encoded the bongkrekic acid biosynthetic gene cluster as shown by the dashed oval and were collectively designated as Group 1 *B. gladioli*. The scale bar represents the number of base substitutions per site.

### Ecological and disease associations of *B. gladioli* evolutionary clades

Given the evolutionary support for the clade restriction of *B. gladioli* pv. *allicola*, and grouping pathovars pv. *gladioli* and pv. *agaricicola* in a separate clade (Figure 2), the ability of selected *B. gladioli* strains to rot eukaryotic tissues was investigated. Mushroom soft-rot bioassays (Roy Chowdhury and Heinemann 2006) demonstrated that *B. gladioli* strains from all 3 phylogenomic groups were capable of decaying mushroom tissue (Figure 3). The assay confirmed the ability of the pv. *agaricicola* reference strain NCPPB 3852 (BCC1883; Figure 3; panel K) to cause disease on its originally associated host. The degree of mushroom rot observed varied, with severe degradation of the mushroom cap tissue most apparent in clade 2 and 3 strains, compared with clade 1 producing less extensive rot (Figure 3). The pathovar *agaricicola-*like strains therefore did not appear specifically adapted to degrade mushroom tissue. *B. gladioli* from all 3 clades also showed conserved plant tissue degradation capabilities within an onion soft-rot model (Jacobs et al 2008). A variable onion rot phenotype was observed for each strain, with the most extensive tissue pitting seen in clades 2 and clade 3 (Figure S1). Overall, rotting capability was demonstrated by the *B. gladioli* strains from all genetic groups.

**Figure 3.**
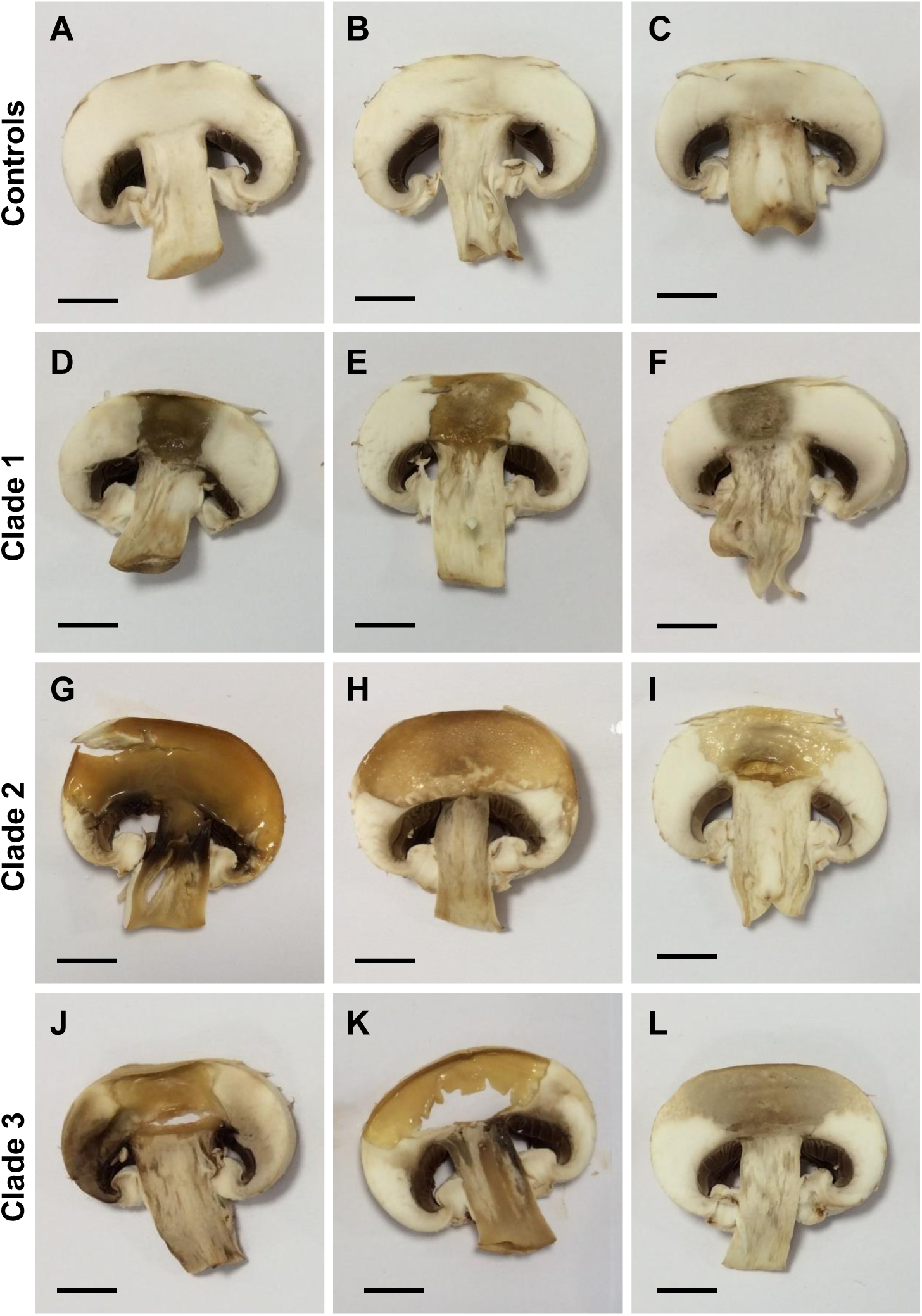
*B. gladioli* from all evolutionary clades are capable of mushroom rot. The ability of *B. gladioli* strains to degrade mushroom tissue was tested by inoculating commercial mushroom tissue slices with standardized bacterial cultures. Inoculated mushrooms were incubated at 30°C for 48 hours with the following shown in each panel (each row are either controls or *B. gladioli* clades as indicated on the left): (A) No treatment control, (B) TSB only, (C) *E. coli* NCTC 12241, (D) *B. gladioli* Clade 1A strain BCC1710, (E) Clade 1B strain BCC1675, (F) Clade 1C strain BCC1678, (G) *B. gladioli* Clade 2 strain BCC1731, (H) *B. gladioli pv. allicola* reference Clade 2 strain BCC1645, (I) B. *gladioli pv. allicola* reference Clade 2 strain BCC1646, (J) *B. gladioli* Clade 3 strain BCC0238, (K) *B. gladioli pv. agaricola* reference Clade 3 strain BCC1883 (NCPPB 3852), (L) *B. gladioli pv. gladioli* reference Clade 3 strain BCC771 (LMG 2216^T^). Pitting and tissue degradation was apparent in all *B. gladioli* inoculated mushrooms; a scale bar (1 cm) is shown in each panel to enable comparison.

Since 94% of the 206 *B. gladioli* strain collection derived from CF lung infections (Table S1), this disease source was also the major origin for each of the evolutionary clades, demonstrating that opportunistic human pathogenicity was also a conserved species phenotype (Figure 2). For the 181 CF strains originating from the United States of America, mapping the state location of the submitting CF treatment centre showed that *B. gladioli* infections were geographically widespread with no phylogeographic linkages to clade types (Figure S2). The Group 1 *B. gladioli* strains (Figure 1 and 2) with the ability to produce bongkrekic acid (see below) were also found to be capable of causing CF lung infections, linking them to opportunistic lung disease for the first time.

### *B. gladioli* possess broad antimicrobial bioactivity

*B. gladioli* is known to produce an array of bioactive specialized metabolites including toxoflavin (Lee et al 2016), bongkrekic acid (Moebius et al 2012), enacyloxins (Ross et al 2014a), caryoynencin (Ross et al 2014b), sinapigladioside (Florez et al 2017), gladiolin (Song et al 2017), and icosalides (Dose et al 2018, Jenner et al 2019). Given this wealth of bioactive products, two *B. gladioli* strains representative of each clade within the species population biology (Figure 2) were screened for antimicrobial activity. Metabolite extracts from the spent agar of *B. gladioli* cultures were examined for their anti-Gram-positive, anti-Gram-negative, and antifungal activity respectively. All 10 strains tested demonstrated activity against *S. aureus*. Only the two isolates from *B. gladioli* clade 1C lacked antifungal activity, while the extracts from the *B. gladioli* clade 1C, clade 2 and clade 3 strains possessed anti-Gram-negative activity (Figure 4**a**). Overall, this antimicrobial activity analysis demonstrated that all the *B. gladioli* strains secreted extractable bioactive compounds, but the quantity and spectrum of bioactivity varied (Figure 4**a**).

**Figure 4.**
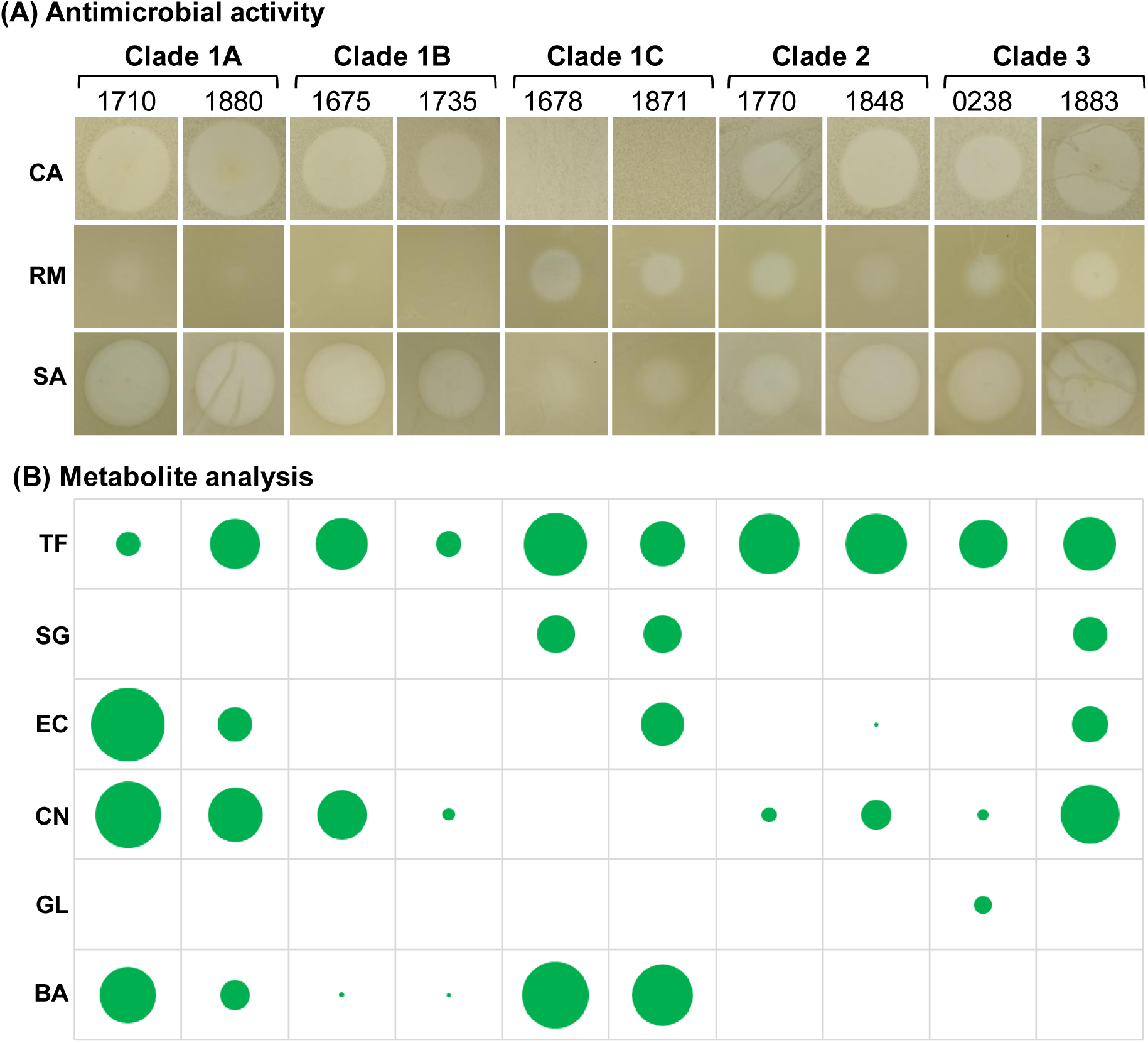
*B. gladioli* antimicrobial activity and bioactive metabolite analysis. **a** The bioactivity of metabolites extracts made from spent growth media from *B. gladioli* cultures. The bioactivity from equivalent metabolite extracts of strains representative of each clade are shown as follows (clade and BCC number are labelled): *B. gladioli* Clade 1A strain BCC1710, Clade 1B strain BCC1675, and Clade 1C strain BCC1678; *B. gladioli* Clade 2 strain BCC1731, *B. gladioli pv. allicola* reference Clade 2 strain BCC1645, and *B. gladioli pv. allicola* reference Clade 2 strain BCC1646; and *B. gladioli* Clade 3 strain BCC0238, *B. gladioli pv. agaricicola* reference Clade 3 strain BCC1883 (NCPPB 3852), and *B. gladioli pv. gladioli* reference Clade 3 strain BCC771 (LMG 2216^T^). Each area of bioactivity was cropped to scale and represents a 3 cm section of the inoculated petri dish. Zones of growth inhibition against *C. albicans* (**CA**), *R. mannitolilytica* (**RM**) and *S. aureus* (**SA**) are labelled as rows. **b** The quantitative analysis of known *B. gladioli* metabolites present in the metabolite extracts as determined by HPLC. Each circle is proportionally scaled to the mean peak height for the following metabolites: toxoflavin (**TF**), sinapigladioside (**SG**), enacyloxin IIa (**EC**), caryoynencin (**CN**), gladiolin (**GL**), and bongkrekic acid (**BA**).

To determine which known metabolites accounted for the *B. gladioli* bioactivity (Figure 4**b**), a combination of HPLC, mass spectrometry and confirmatory BGC pathway mutagenesis was applied. Under the specialized metabolite inducing growth conditions used (Mahenthiralingam et al 2011), toxoflavin was found to be produced by all *B. gladioli* strains (Figure 4**b**). The isothiocyanate sinapigladioside (Florez et al 2017) was produced by both clade 1C strains and one clade 3 strain. Enacyloxin IIa (Mahenthiralingam et al 2011) was present in both clade 1A strains, a clade 1C strain, a clade 3 strain, and was also detected at low quantities within the clade 2 strain BCC1848 (Figure 4**b**). Production of the polyyne caryoynencin (Ross et al 2014b) was widespread and detected in eight of the 10 strains tested, with an absence in the clade 1C strains (Figure 4**b**). Gladiolin (Song et al 2017) was detected in its discovery strain *B. gladioli* BCC0238, but the macrolide was not present in other strains. Bongkrekic acid was detected in all 6 clade 1 strains, although only limited amounts were present in the two clade 1B strains examined (Figure 4**b**). Overall, the metabolite analysis showed that individual *B. gladioli* strains were capable of producing up to 4 different bioactive metabolites (Figure 4**b**) underpinning the broad spectrum of antimicrobial activity of *B. gladioli* (Figure 4**a**).

### Distribution of known specialized metabolite BGCs in *B. gladioli*

To expose the genetic basis of bioactivity and the production of multiple metabolites (Figure 4), the distribution of known metabolite BGCs was mapped by genome mining. Using sequence read mapping to known *B. gladioli* BGCs the presence of toxoflavin, caryoynencin, bongrekic acid, enacyloxin, gladiolin, and icosalide BGCs were determined. Across the phylogenomically defined *B. gladioli* clades, random distribution as well as clade-specific presence of BGCs was observed (Figure 5**a**). The capacity for toxoflavin, caryoynencin and icosalide biosynthesis was widely distributed across *B. gladioli*, with the toxoflavin BGC being absent in only two of the 206 strains. The caryoynencin BGC was uniquely absent in all 10 clade 1C strains (correlating to a lack of detection of the metabolite; Figure 4**b**). The icosalide BGC mirrored this clade 1C absence, but also showed random loss in six other strains from across *B. gladioli* (one clade 1B, four clade 2 and one clade 3 strains; Figure 5**a**). The gladiolin and bongkrekic BGCs demonstrated evolutionary restrictions to specific clades as follows. A total of 83 of the 106 clade 3 strains (78%) encoded the gladiolin BGC and it was absent from the other *B. gladioli* clades. All 27 strains within clades 1A, 1B and 1C, encoded the BGC for bongkrekic acid production, validating its presence as marker to collectively designate them as Group 1 *B. gladioli* strains (Figure 1), despite their distinct nature as evolutionary clades (Figure 2).

**Figure 5.**
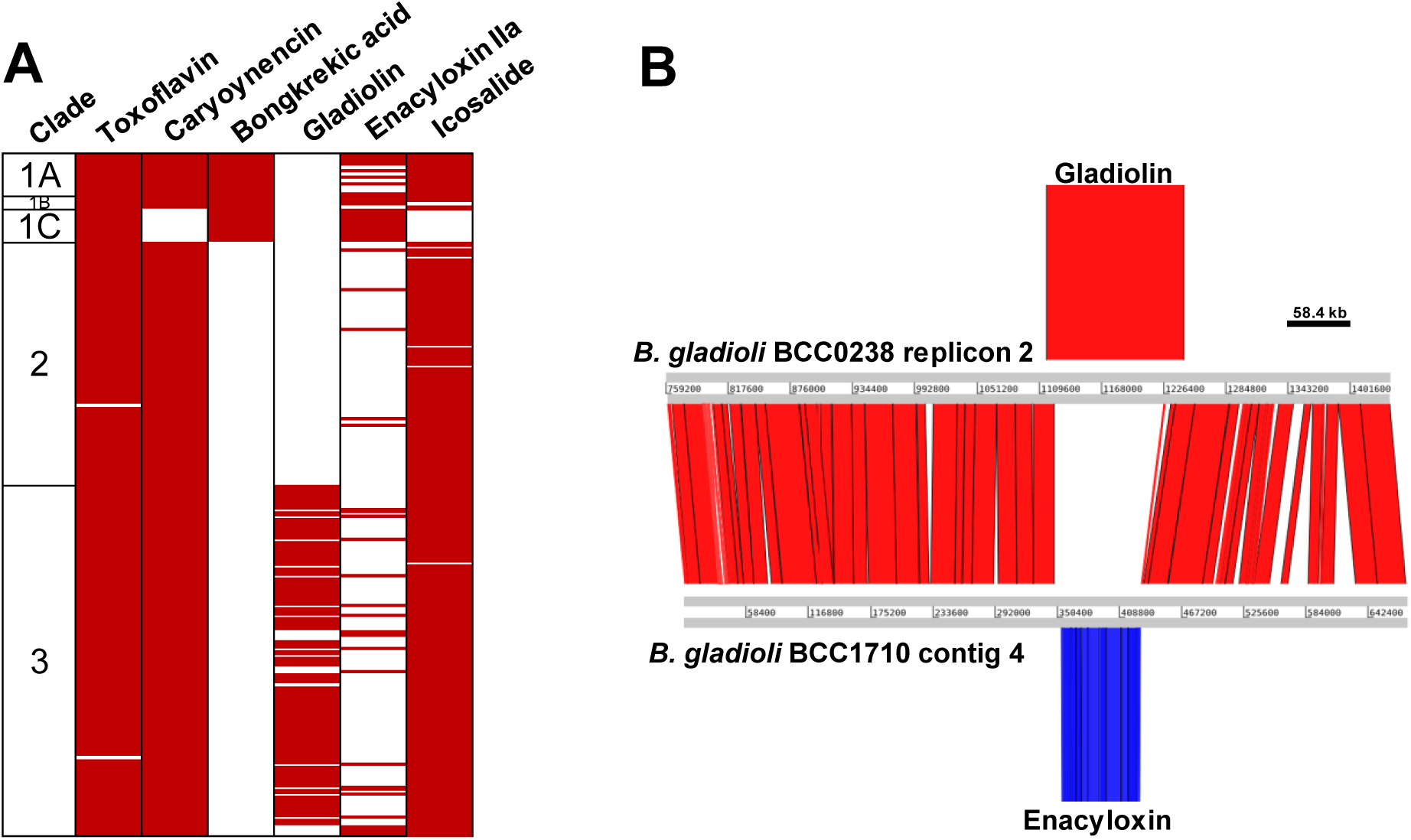
Distribution of known *B. gladioli* specialized metabolite BGCs and common genomic location for gladiolin and enacyloxin biosynthesis. **a** Sequence reads were mapped to known specialized metabolite BGCs to determine their presence or absence within the 206 the *B. gladioli* genomes. BGC presence indicated by red shading and columns from left to right show the *B. gladioli* clade, and BGCs for toxoflavin, caryoynencin, bongkrekic acid, gladiolin, enacyloxin IIa and icosalide. **b** A genomic comparison plot was constructed using the Artemis Comparison Tool for *B. gladioli* BCC0238 (clade 3) and BCC1710 (clade 1A). The common insertion point for either the gladiolin or enacyloxin BGCs in these strains is shown, together with extensive genomic synteny upstream and downstream of this specialized metabolite encoding location (scale bar indicates 58.4 kb).

The enacyloxin BGC was randomly distributed within *B. gladioli* (Figure 5**a**). Its presence within clade 1 strains was most conserved with 21 of 27 strains (77%) being enacyloxin BGC positive and 100% of clade 1C strains encoding it. Interestingly, no strain within the collection of 206 encoded both the enacyloxin and gladiolin BGC (Figure 5**a**). Genomic interrogation of this inverse correlation led to the discovery that these large polyketide BGCs occupied the same genetic locus within *B. gladioli* when present (Figure 5**b**). This conserved region of the genome was on the second genomic replicon of *B. gladioli* and either encoded enacyloxin (43 strains) or gladiolin (83 strains) or remained empty in terms of specialized metabolite BGCs (80 strains). Upstream and downstream of these polyketide BGC island insertion points were blocks of conserved and syntenic genomic DNA. These surrounding regions of the *B. gladioli* second genomic replicon did not show the presence of mobile DNA markers to indicate the BGC insertion point acted as a gene capture hotspot.

### *B. gladioli* bongkrekic acid biosynthesis: a new potential risk factor for CF lung infection

In total, 25 of the 27 strains within clades 1A, 1B and 1C, were recovered from CF infection (Table S1 and Figure 2), and all possessed the bongkrekic acid BGC (Figure 5A). To date this lethal toxin has only been associated with food poisoning related *B. gladioli* (Gudo et al 2018, Jiao et al 2003, Moebius et al 2012) and not linked to disease in people with CF. Analysis of 12 toxin BGC positive strains showed that 11 of them produced bongkrekic acid *in vitro*, but the extent of toxin production by each was variable (Figure S4). Four of these bongkrekic acid producers (BCC1675, BCC1686, BCC1701 and BCC1710; Figure S4) were subsequently grown in artificial CF sputum medium (Kirchner et al 2012), and with the exception of strain BCC1701 (a low producer; Figure S4), toxin production was detected in the remaining 3 strains by LC-MS. This toxin production under CF lung infection-like growth conditions, prompted the development of diagnostic PCR probes to enable rapid identification of the bongkrekic acid positive *B. gladioli* as a potential clinical risk marker for CF. Testing of this *bonA* gene (Moebius et al 2012) targeting PCR (Figure S6) on 122 of the *B. gladioli* CF strains prior to their genome sequencing identified 13 isolates as positive. Subsequent genome sequencing demonstrated that all encoded a complete BGC for the toxin (Figure 5**a**) validating the risk marker PCR.

Since CF patients are administered multiple antibiotics to suppress their lung infections. A recent finding that low concentrations of antibiotics may act as inducers of *Burkholderia* specialized metabolites (Seyedsayamdost 2014) added further potential risk to the occurrence of bongkrekic toxin positive *B. gladioli* strains within CF individuals. Furthermore, the antibiotic trimethoprim, which is widely used for treatment of *Burkholderia* CF infections, was specifically shown to be a highly effective elicitor of specialized metabolite production in *B. thailandensis* (Okada et al 2016), compounding the threat of toxin activation. To test antibiotic induced BGC expression (Okada et al 2016, Seyedsayamdost 2014), six *B. gladioli* CF strains possessing a range of bongkrekic acid production levels (Figure 6**a**) were screened for toxin production with and without the presence of sub-inhibitory levels of trimethoprim. In contrast to the *B. thailandensis* metabolite activation (Okada et al 2016), no *B. gladioli* strains demonstrated induction of bongkrekic acid by exposure to trimethoprim, but five of the six strains analysed showed a clear suppression of toxin production (*B. gladioli* BCC1678, Figure 6**b** and 6**c**; Figure S6 shows the data for all six strains tested).

**Figure 6.**
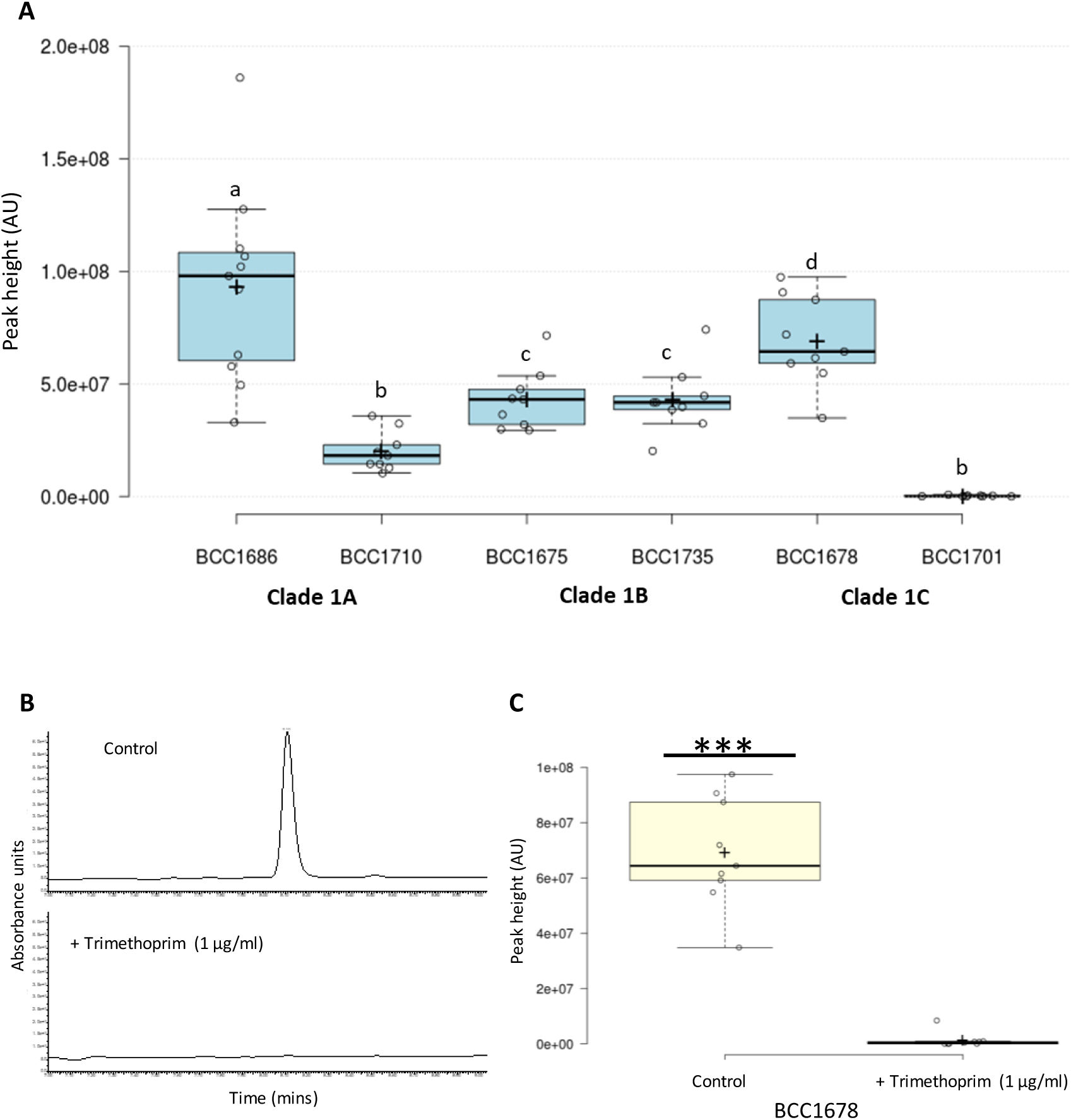
Subinhibitory concentrations of trimethoprim reduce bongkrekic acid production by *B. gladioli* clade 1 strains. **a** HPLC analysis of 6 *B. gladioli* strains (strain numbers are shown) belonging to clades 1A, 1B and 1C demonstrated variable levels of bongkrekic acid toxin production by HPLC analysis of their metabolite extracts (peak height is plotted). Differences between the mean values of bongkrekic acid were determined using the LSD test α=0.05. Means followed by the same letter are not significantly different. **b** The presence of subinhibitory concentrations of trimethoprim (1 µg/ml) reduced the production of bongkrekic acid in strain BCC1678 as shown the HPLC metabolite analysis comparing production levels against the control condition without antibiotic. **c** Quantitative comparison of bongkrekic acid production by strain BCC1678 in the presence and absence of trimethoprim (n=9) shows the significant (p<0.001) suppression of toxin biosynthesis caused by antibiotic exposure. Statistical significance was determined using a two-tailed t-test.

## DISCUSSION

In the last decade the specialized metabolites produced by *Burkholderia* have been extensively studied and multiple compounds have been shown to be activated or functional in different ecological settings (Kunakom and Eustaquio 2019). *B. gladioli* shows a very wide range of beneficial (Coenye et al 2000, Florez et al 2017, Song et al 2017) versus detrimental traits (Coenye et al 1999, Gudo et al 2018, Hildebrand et al 1973, Jiao et al 2003, Quon et al 2011), several of which relate to specialized metabolite production. With multiple taxonomic re-classifications (Coenye et al 1999, Coenye et al 2000), an unknown basis for pathovar status in plant disease (Coenye et al 1999, Hildebrand et al 1973, Wright et al 1993), calls for a specific recognition of the lethal food-poisoning “*cocovenenans*” pathovar (Jiao et al 2003), an emerging presence in CF lung infection (Lipuma 2010), and its expanding role as a source of specialized metabolites (Dose et al 2018, Jenner et al 2019, Kunakom and Eustaquio 2019, Ross et al 2014a, Ross et al 2014b, Song et al 2017), there was a clear need to understand the population ecology of *B. gladioli*.

The ability of individual strains of *B. gladioli* to produce both beneficial or toxic specialized metabolites is also very interesting given their wide interactions with different eukaryotic hosts (Florez et al 2017, Lee et al 2016). Based on the ANI-genomic species concept (Goris et al 2007), our novel genomic analyses confirmed that all pathovars of *B. gladioli* collectively comprise a single species. However, from the phylogenomic analysis, we have uniquely shown that *B. gladioli* comprises 5 evolutionarily distinct clades (Figure 2). The capacity to encode and produce the bongkrekic acid toxin was specific to 3 of these clades (1A, 1B and 1C) and provided a unifying, and potentially medically relevant feature to designate them as a single group. These bongkrekic acid positive strains were also linked for the first time as a potential risk factor for people with CF. While production of the toxin during infection and its association with poor disease outcome needs to be determined, we have showed that toxin BGC is biosynthetically active in Group 1 *B. gladioli* strains recovered from people with CF (Figure S4).

The population biology and pathogenicity traits of very few *Burkholderia* species has been described in depth using phylogenomics. The transmission dynamics of a *Burkholderia dolosa* CF outbreak was tracked in 14 patients over 16 years and identified mutations in the 112 isolates that showed parallel evolution towards increased antibiotic resistance and tolerance of low oxygen (Lieberman et al 2011). The cause of melioidosis, *Burkholderia pseudomallei*, has been subjected to arguably the most extensive genomic characterisation because of its pathogenicity and due to its threat as a bioterrorism agent (Chewapreecha et al 2017). Genome sequencing of 469 *B. pseudomallei* isolates showed the species comprised two distinct populations, an ancestral Australian reservoir that anthropogenically transmitted and diverged within Asia, and spread further via the slave trade from Africa to South America (Chewapreecha et al 2017). Mapping the phylogenomics of *B. ambifaria* as a historically used biopesticide revealed the BGC for cepacin A, a key antimicrobial mediating plant protection against pathogenic oomycetes (Mullins et al 2019). However, the population biology of plant pathogenic *Burkholderia* has not been studied in depth and our genomic analysis of *B. gladioli* is unique in uncovering whether plant pathovar status has an evolutionary basis. Only the *B. gladioli* pv. *allicola* associated with onion soft-rot plant disease were evolutionarily distinct as clade 2 strains (Figure 2), but rotting capability was associated with all clades and historical pathovars (Figure 3 and Figure S1). The specific genetic factors linked to the separation of pathovar *allicola* as clade 2 and its distinction from clade 3 plant disease strains remain to be determined.

In contrast to the broad conservation of plant disease traits across *B. gladioli*, bongkrekic acid producing strains (Gudo et al 2018, Jiao et al 2003, Moebius et al 2012) associated with fatal human food poisoning were more closely related. They were designated Group 1 by their ANI relatedness (Figure 1) and unifying bongkrekic acid BGC presence (Figure 5). Although the human disease Group 1 *B. gladioli* pv. *cocovenenans* strains were not a distinct species (Figure 1) and occupied 3 distinct evolutionary clades (Figure 2), this 100% bongkrekic acid BGC positivity adds weight to the call for their differentiation as a distinct toxin-positive *B. gladioli* subgroup (Jiao et al 2003). With 13% of the *B. gladioli* CF isolates examined encoding the bongkrekic acid BGC, this is a worrying potential risk factor for *Burkholderia* lung infections, especially since toxin production can occur under lung infection-like growth conditions, such as artificial CF sputum. The clinical outcome of *Burkholderia* infection is frequently highly variable (Frangolias et al 1999) and severe systemic disease has been associated with *B. gladioli* in CF (Jones et al 2001). With the ability to rapidly identify bongkrekic acid positive *B. gladioli* using a PCR diagnostic (Figure S5), we are now in a strong position to understand if the toxin plays a role in poor clinical outcome of infected CF patients. Also, since we have shown that trimethoprim acts to suppress toxin production (Figure 6), rather than activate this *Burkholderia* specialized metabolite (Kirchner et al 2012), a case can be made for antibiotic therapy to be maintained in bongkrekic acid positive *B. gladioli* CF infection.

By combining genomics with analytical chemistry, we have also been able to map the repertoire of bioactive specialized metabolite BGCs across the *B. gladioli* as a species. This demonstrated that the bioactivity of *B. gladioli* is frequently the result of the production of multiple metabolites (Figure 4). The widespread distribution and conservation of BGCs for toxoflavin (Lee et al 2016), caryoynencin (Ross et al 2014b) and the icosalides (Dose et al 2018, Jenner et al 2019) suggest they are ancestral to *B. gladioli* as a species (Figure 5). The specificity of gladiolin biosynthesis to clade 3 strains also sheds light on the classification of the recently identified symbiotic *B. gladioli* strains that protect the eggs of Lagriinae beetles from fungal attack (Florez et al 2017). The symbiotic beetle *B. gladioli* strain Lv-StA encodes the BGC for the antibiotic lagriene (Florez et al 2017) which identical to the BGC for the macrolide gladiolin (Song et al 2017). Since the gladiolin BGC, and by default the lagriene BGC, is restricted to *B. gladioli* clade 3, the insect symbionts must be members of this clade. It is also clear that all *B. gladioli* clades are geographically widely distributed from the analysis of US CF infection strains (Figure S2). The ecological significance of herbivorous Lagriinae and other beetles in distributing the such bacterial symbionts across continental ranges will be fascinating to understand.

Within this study, we were able to gain an insight into the ecological distribution of *B. gladioli* by sampling the opportunistic infections the bacterium causes in people with CF. In the absence of patient-to-patient or common source transmission, the natural environment is the main source of *Burkholderia* CF lung infections (Lipuma 2010). From the US CF patient data (Figure S2), all *B. gladioli* clades appear widely distributed across a continental range. Soil, the rhizosphere and terrestrial freshwater environments are common sources of *Burkholderia* (Suarez-Moreno et al 2012). Outside of CF infection (Lipuma 2010), plant disease (Coenye et al 1999, Hildebrand et al 1973, Wright et al 1993) or food-poisoning (Jiao et al 2003), little is known about other sources of *B. gladioli*. Recent findings of close associations with insects (Florez et al 2017) and fungi (Dose et al 2018, Jenner et al 2019) point to multiple symbiotic roles *B. gladioli* may have in the natural environment. The population biology, pathogenicity, metabolite and BGC analysis we have carried out provides a systematic framework upon which the ecological distribution of *B. gladioli* can be mapped.

## ACKNOWLEDGEMENTS

This research was supported by a grant from the BBSRC (BB/L021692/1) to E.M, G.L.C, T.R.C and J.P; A.J.M., G.W. and E.M also acknowledged current funding from grant BB/S007652/1. E.M. and G.W. acknowledge funding from the Life Science Bridging Fund LSBF R2-004 for the HPLC analysis and instrumentation. T.R.C and M.J.B acknowledge funding from the MRC (MR/L015080/1). The Bruker MaXis II instrument used in this study was funded by the BBSRC (BB/M017982/1). M.J. is supported by a BBSRC Future Leader Fellowship (BB/R012121/1) and G.L.C. is the recipient of a Wolfson Research Merit Award from the Royal Society (WM130033).

## AUTHOR CONTRIBUTIONS

(CRediT taxonomy; https://casrai.org/credit/)

We describe author contributions to the paper using the CRedit taxonomy. *Conceptualisation:* E.M, and C.J.; *Data Curation:* E.M., C.J., G.W., A.J.M., M.J., J.P., J.J.L., and G.L.C.; *Formal analysis:* E.M., C.J., A.J.M., M.J., Y.D., J.P., J.J.L., and G.L.C; *Funding Acquisition:* E.M., J.P., G.LC. and J.J.L.; *Investigation:* E.M., C.J., G.W., A.J.M., M.J., J.P., T.S., J.J.L., and G.L.C; *Methodology:* C.J., G.W., A.J.M., M.J., Y.D., M.J.B., and T.R.C.; *Project Administration:* E.M., J.P. and G.L.C; *Resources:* E.M., J.P., J.J.L., and G.L.C.; *Software:* C.J., A.J.M., M.J.B., and T.R.C.; *Supervision:* E.M., J.P. and G.L.C.; *Validation:* E.M., C.J; *Visualisation:* C.J., G.W. and E.M.; *Writing-Original Draft:* E.M. and C.J.; *Writing-Review & Editing:* all authors.

