## Supplementary information file for "Kill and cure: genomic phylogeny and bioactivity of a diverse collection of *Burkholderia gladioli* bacteria capable of pathogenic and beneficial lifestyles"

[8] Current address: Department of Veterinary Medicine, Madingley Road, Cambridge, CB3 0ES

**Running title:** Population genomics of *B. gladioli*

**Key words:** *Burkholderia, B. gladioli*, antibiotic production, plant pathogenesis, cystic fibrosis infection, phylogenomics.

**CONTENTS:**

**SUPPLEMENTARY TABLES**

**Table S1.** Collection of *Burkholderia gladioli* isolates assembled and analysed in this study

**Table S2**. Primers and *B. gladioli* isolates used for generation of biosynthetic gene cluster insertional mutants

**SUPPLEMENTARY FIGURES**

**Figure S1.** *B. gladioli* from all evolutionary clades are capable of onion rot

**Figure S2**. CF infection associated with each *B. gladioli* clade is widely distributed across the United States of America

**Figure S3.** UHPLC-ESI-Q-TOF-MS analysis of metabolites produced by *B. gladioli* isolates

**Figure S4.** *B. gladioli* clade 1 strains have a variable ability to produce bongkrekic acid

**Figure S5.** Detection of *B. gladioli* clade 1 strains by bongkrekic acid *bonA* gene PCR

**Figure S6.** Subinhibitory trimethoprim reduces bongrekic acid production

### Table S1: Collection of *Burkholderia gladioli* isolates assembled and analysed in this study

| Strain Name | Phylogenomic Clade | **Strain Details** (CF = CF; ENV = environmental, ENVI = environmental from industry) | Illumina Sequencing | | | | | Analysis Performed | | | | | | |
|---|---|---|---|---|---|---|---|---|---|---|---|---|---|---|
| | | | Accession (PacBio) | Genome Size | Number of Contigs | %GC | N50 | MLST sequence type | Core Genome Analysis | Mushroom Rot | Bioactivity Screening | HPLC Analysis | BGC Mapping | Bongkrekic acid analysis |
| BCC1650 | 1A | ENV; *B. gladioli* pv *cocovenenans*, LMG 11626, Indonesia | ERS784828 | 8091943 | 35 | 68.24 | 708245 | 957 | | | | BCC1650 | | |
| BCC1661 | 1A | CF Isolate, USA | ERS784861 | 8327590 | 45 | 68.06 | 632785 | 959 | | | | | | |
| BCC1665 | 1A | CF Isolate, USA | ERS784925 | 8328710 | 45 | 68.06 | 601921 | 963 | | | | | | |
| BCC1675 | 1B | CF Isolate, USA | ERS785062 | 8080953 | 47 | 68.09 | 936495 | 970 | | BCC1675 | BCC1675 | BCC1675 | | BCC1675 |
| BCC1678 | 1C | CF Isolate, USA | ERS784880 | 7819531 | 96 | 68.04 | 235084 | 973 | | BCC1678 | BCC1678 | BCC1678 | | BCC1678 |
| BCC1686 | 1A | CF Isolate, USA | ERS784816 | 8334290 | 38 | 68.06 | 728975 | 963 | | | | | | BCC1686 |
| BCC1689 | 1A | CF Isolate, USA | ERS784865 | 8252657 | 47 | 67.94 | 353209 | 993 | | | | | | |
| BCC1692 | 1A | CF Isolate, USA | ERS784914 | 8387798 | 41 | 67.98 | 810454 | 983 | | | | | | |
| BCC1697 | 1A | CF Isolate, USA | ERS784984 | 8085377 | 49 | 68.02 | 652756 | 989 | | | | | | |
| BCC1701 | 1C | CF Isolate, USA | ERS784867 | 8145614 | 94 | 67.98 | 280973 | 993 | | | | BCC1701 | | BCC1701 |
| BCC1710 | 1A | CF Isolate, USA | ERS785039 (GCA_900608535) | 8374934 | 44 | 68.02 | 615977 | 999 | | BCC1710 | BCC1710 | BCC1710 | | BCC1710 |
| BCC1735 | 1B | CF Isolate, USA | ERS785008 | 7922226 | 64 | 67.98 | 388060 | 1019 | | | BCC1735 | BCC1735 | | BCC1735 |
| BCC1766 | 1A | CF Isolate, USA | ERS1371589 | 8275704 | 144 | 67.89 | 275070 | Novel | | | | | | |
| BCC1780 | 1B | CF Isolate, USA | ERS1328777 | 8428886 | 208 | 67.76 | 220441 | Novel | | | | | | |
| BCC1781 | 1C | CF Isolate, USA | ERS1328887 | 7974402 | 195 | 68.16 | 208905 | Novel | | | | | | |
| BCC1812 | 1A | CF Isolate, USA | ERS1328895 | 8211259 | 162 | 67.94 | 301688 | Novel | | | | | | |
| BCC1819 | 1A | CF Isolate, USA | ERS1328796 | 8410357 | 115 | 67.9 | 331109 | Novel | | | | | | |
| BCC1821 | 1B | CF Isolate, USA | ERS1328797 | 8122457 | 126 | 68.08 | 263017 | Novel | | | | | | |
| BCC1829 | 1A | CF Isolate, USA | ERS1371601 | 8600125 | 112 | 67.8 | 317992 | Novel | | | | | | |
| BCC1837 | 1C | CF Isolate, USA | ERS1371606 | 8128598 | 159 | 67.99 | 206652 | Novel | | | | | | |
| BCC1843 | 1C | CF Isolate, USA | ERS1328808 | 7992663 | 209 | 68.1 | 177545 | Novel | | | | | | |
| BCC1844 | 1C | CF Isolate, USA | ERS1328903 | 8186135 | 206 | 67.98 | 219436 | Novel | | | | | | |
| BCC1861 | 1C | CF Isolate, USA | ERS1371619 | 8609383 | 249 | 67.75 | 216674 | Novel | | | | | | |
| BCC1864 | 1C | CF Isolate, USA | ERS1328908 | 8490658 | 220 | 67.9 | 212901 | Novel | | | | | | |

| | | | | | | | | | | | | |
|---|---|---|---|---|---|---|---|---|---|---|---|---|
| BCC1870 | 1C | CF Isolate, USA | ERS1371625 | 8341408 | 182 | 67.95 | 186380 | Novel | | | | |
| BCC1871 | 1C | CF Isolate, USA | ERS1371626 | 8043875 | 155 | 68.11 | 284538 | Novel | | | BCC1871 | BCC1871 |
| BCC1880 | 1A | ENV; *B. gladioli* pv *cocovenenans*, LMG 18113, China | ERS1371628 | 8222883 | 77 | 68.04 | 621809 | Novel | | | | |
| BCC0252 | 2 | CF Isolate, Australia | ERS784955 | 8212491 | 63 | 68.09 | 289654 | 947 | | | | |
| BCC1619 | 2 | CF Isolate, USA | ERS784817 | 7880197 | 36 | 68.13 | 868935 | 554 | | | | |
| BCC1620 | 2 | CF Isolate, USA | ERS784832 | 8184546 | 61 | 68.09 | 416357 | 951 | | | | |
| BCC1621 | 2 | CF Isolate, USA | ERS784849 (GCA_900608525) | 8467292 | 76 | 67.95 | 434143 | 903 | | | | |
| BCC1624 | 2 | CF Isolate, USA | ERS784895 | 7876116 | 48 | 68.14 | 430243 | 554 | | | | |
| BCC1645 | 2 | ENV; *B. gladioli* pv. *alliicola*, LMG 6954, Australia | ERS784911 | 8603880 | 78 | 67.87 | 383325 | 903 | | | | BCC1645 |
| BCC1646 | 2 | ENV; *B. gladioli* pv. *alliicola*, LMG 6878, India | ERS784928 | 8379242 | 87 | 67.96 | 286552 | 954 | | | | |
| BCC1648 | 2 | ENV; *B. gladioli* pv. *alliicola*, LMG 2121, USA | ERS784957 | 8607718 | 85 | 67.87 | 310631 | 903 | | | | |
| BCC1662 | 2 | CF Isolate, USA | ERS784878 | 8507695 | 67 | 68.05 | 410006 | 947 | | | | |
| BCC1663 | 2 | CF Isolate, USA | ERS784894 | 8510108 | 64 | 68.05 | 410015 | 947 | | | | |
| BCC1664 | 2 | CF Isolate, USA | ERS784910 | 8498792 | 73 | 68.06 | 378407 | 947 | | | | |
| BCC1666 | 2 | CF Isolate, USA | ERS784942 | 8506933 | 67 | 68.05 | 408038 | 947 | | | | |
| BCC1667 | 2 | CF Isolate, UK | ERS784960 | 8480007 | 78 | 67.84 | 427447 | 965 | | | | |
| BCC1670 | 2 | CF Isolate, UK | ERS784814 | 8282986 | 75 | 68.03 | 309426 | 903 | | | | |
| BCC1671 | 2 | CF Isolate, UK | ERS784830 | 8347055 | 76 | 68.09 | 369886 | 968 | | | | |
| BCC1672 | 2 | CF Isolate, UK | ERS784846 | 8272557 | 75 | 68.03 | 287015 | 903 | | | | |
| BCC1674 | 2 | CF Isolate, UK | ERS784994 | 8431622 | 84 | 67.87 | 356821 | 965 | | | | |
| BCC1676 | 2 | CF Isolate, USA | ERS784863 | 7987029 | 52 | 68.26 | 467101 | 971 | | | | |
| BCC1679 | 2 | CF Isolate, USA | ERS784896 | 8113442 | 62 | 68.09 | 272665 | 971 | | | | |
| BCC1680 | 2 | CF Isolate, USA | ERS784912 | 7965535 | 52 | 68 | 411881 | 554 | | | | |
| BCC1681 | 2 | CF Isolate, USA | ERS784927 | 7321015 | 56 | 68.09 | 382283 | 975 | | | | BCC1681 |
| BCC1682 | 2 | CF Isolate, USA | ERS784944 | 7942355 | 57 | 68.03 | 428648 | 947 | | | | |
| BCC1684 | 2 | CF Isolate, USA | ERS784975 | 8383929 | 56 | 67.96 | 576064 | 971 | | | | |
| BCC1685 | 2 | CF Isolate, USA | ERS784983 | 7943459 | 66 | 68.03 | 314381 | 947 | | | | |
| BCC1690 | 2 | CF Isolate, USA | ERS784881 | 8470418 | 75 | 67.94 | 433041 | 903 | | | | |
| BCC1693 | 2 | CF Isolate, USA | ERS784929 | 8210869 | 47 | 68.1 | 668608 | 554 | | | | |
| BCC1695 | 2 | CF Isolate, USA | ERS784964 | 7905325 | 53 | 68.06 | 538303 | 971 | | | | |

| | | | | | | | | | | | | |
|---|---|---|---|---|---|---|---|---|---|---|---|---|
| BCC1699 | 2 | CF Isolate, USA | ERS784834 | 8345011 | 78 | 67.87 | 354820 | 991 | | | | |
| BCC1705 | 2 | CF Isolate, USA | ERS784916 | 8145336 | 49 | 68.17 | 551683 | 995 | | | | |
| BCC1708 | 2 | CF Isolate, USA | ERS784966 | 8331226 | 49 | 68.03 | 945197 | 554 | | | | |
| BCC1711 | 2 | CF Isolate, USA | ERS785006 | 8344918 | 67 | 68.06 | 335836 | 903 | | | | |
| BCC1712 | 2 | CF Isolate, USA | ERS785025 | 8201991 | 62 | 68.08 | 388388 | 971 | | | | |
| BCC1715 | 2 | CF Isolate, USA | ERS784977 | 7963860 | 43 | 68.14 | 954999 | 1003 | | | | |
| BCC1716 | 2 | CF Isolate, USA | ERS785052 | 8495662 | 77 | 67.87 | 385075 | 554 | | | | |
| BCC1717 | 2 | CF Isolate, USA | ERS785014 | 8418479 | 69 | 67.93 | 430020 | 1004 | | | | |
| BCC1719 | 2 | CF Isolate, USA | ERS785042 | 7988187 | 68 | 68.17 | 429154 | 903 | | | | |
| BCC1731 | 2 | CF Isolate, USA | ERS784997 | 8485637 | 96 | 67.97 | 339104 | 1016 | | BCC1731 | | |
| BCC1756 | 2 | CF Isolate, USA | ERS1328753 | 7729600 | 167 | 68.11 | 264066 | 903 | | | | |
| BCC1759 | 2 | CF Isolate, USA | ERS1333276 | 8389900 | 206 | 67.93 | 202663 | Novel | | | | |
| BCC1761 | 2 | CF Isolate, USA | ERS1371587 | 8105698 | 114 | 68.15 | 227067 | Novel | | | | |
| BCC1762 | 2 | CF Isolate, USA | ERS1328756 | 8362320 | 214 | 67.86 | 232045 | Novel | | | | |
| BCC1764 | 2 | CF Isolate, USA | ERS1371588 | 7946735 | 117 | 68.19 | 273951 | Novel | | | | |
| BCC1768 | 2 | CF Isolate, USA | ERS1371591 | 8251454 | 139 | 68.11 | 226248 | Novel | | | | |
| BCC1769 | 2 | CF Isolate, USA | ERS1328884 | 8343545 | 225 | 68.02 | 248528 | 554 | | | | |
| BCC1770 | 2 | CF Isolate, USA | ERS1328772 | 8405384 | 148 | 67.98 | 261993 | 554 | | | BCC1770 | BCC1770 |
| BCC1771 | 2 | CF Isolate, USA | ERS1328943 | 8398051 | 128 | 67.99 | 303762 | 554 | | | | |
| BCC1773 | 2 | CF Isolate, USA | ERS1328885 | 8072772 | 131 | 68.21 | 264067 | 903 | | | | |
| BCC1774 | 2 | CF Isolate, USA | ERS1371592 | 7885091 | 119 | 68.1 | 264045 | 903 | | | | |
| BCC1779 | 2 | CF Isolate, USA | ERS1328944 | 8227602 | 296 | 68.02 | 190616 | 903 | | | | |
| BCC1782 | 2 | CF Isolate, USA | ERS1328778 | 7984599 | 133 | 68.24 | 264129 | 971 | | | | |
| BCC1783 | 2 | CF Isolate, USA | ERS1336060 | 8376932 | 184 | 67.96 | 224024 | 965 | | | | |
| BCC1785 | 2 | CF Isolate, USA | ERS1328779 | 7979574 | 139 | 68.26 | 243147 | 971 | | | | |
| BCC1790 | 2 | CF Isolate, USA | ERS1333278 | 8396898 | 197 | 67.9 | 249452 | 903 | | | | |
| BCC1792 | 2 | CF Isolate, USA | ERS1328890 | 8115078 | 129 | 68.17 | 276382 | 1508 | | | | |
| BCC1795 | 2 | CF Isolate, USA | ERS1328784 | 7813137 | 123 | 68.17 | 275050 | Novel | | | | |
| BCC1798 | 2 | CF Isolate, USA | ERS1371596 | 8241429 | 151 | 68.07 | 230356 | 971 | | | | |
| BCC1803 | 2 | CF Isolate, USA | ERS1371597 | 7705345 | 124 | 68.12 | 272468 | 554 | | | | |
| BCC1804 | 2 | CF Isolate, USA | ERS1371598 | 8439232 | 162 | 68 | 250984 | 971 | | | | |
| BCC1805 | 2 | CF Isolate, USA | ERS1328789 | 7850130 | 140 | 67.94 | 270833 | 554 | | | | |

| | | | | | | | | | | | | |
|---|---|---|---|---|---|---|---|---|---|---|---|---|
| BCC1814 | 2 | CF Isolate, USA | ERS1336062 | 8201270 | 156 | 68.14 | 206859 | 975 | | | | |
| BCC1815 | 2 | CF Isolate, USA | ERS1328794 | 8940443 | 186 | 67.39 | 262346 | 954 | | | | |
| BCC1820 | 2 | CF Isolate, USA | ERS1328897 | 8038288 | 131 | 68.15 | 242112 | Novel | | | | |
| BCC1825 | 2 | CF Isolate, USA | ERS1328799 | 8200736 | 119 | 67.96 | 211609 | 971 | | | | |
| BCC1828 | 2 | CF Isolate, Canada | ERS1328899 | 8234597 | 153 | 68.04 | 268035 | 971 | | | | |
| BCC1831 | 2 | CF Isolate, Canada | ERS1371603 | 8416509 | 153 | 68 | 189469 | 947 | | | | |
| BCC1832 | 2 | CF Isolate, USA | ERS1371604 | 8156592 | 135 | 68.08 | 208866 | 971 | | | | |
| BCC1834 | 2 | CF Isolate, USA | ERS1328950 | 7897744 | 118 | 68.06 | 286322 | 991 | | | | |
| BCC1835 | 2 | CF Isolate, USA | ERS1371605 | 8392649 | 159 | 67.96 | 264845 | 554 | | | | |
| BCC1838 | 2 | CF Isolate, USA | ERS1336064 | 8426173 | 204 | 67.91 | 179624 | 947 | | | | |
| BCC1848 | 2 | CF Isolate, USA | ERS1328904 | 8130201 | 112 | 68.04 | 301164 | Novel | | | BCC1848 | BCC1848 |
| BCC1855 | 2 | CF Isolate, USA | ERS1371614 | 8309427 | 177 | 68.01 | 186469 | Novel | | | | |
| BCC1865 | 2 | CF Isolate, USA | ERS1328819 | 8217507 | 131 | 68.15 | 233312 | 971 | | | | |
| BCC1867 | 2 | CF Isolate, USA | ERS1371623 | 8391961 | 158 | 67.96 | 178703 | Novel | | | | |
| BCC0238 | 3 | CF Isolate, Canada | ERS784907 (GCA_900631635.1) | 8541552 | 79 | 68 | 479899 | 946 | | BCC0238 | BCC0238 | BCC0238 |
| BCC0507 | 3 | CF Isolate, Italy | ERS784970 | 8350939 | 61 | 68.13 | 335266 | 948 | | | | |
| BCC0771 | 3 | *B. gladioli* pv. *gladioli*, LMG 2216 (ATCC 10248), USA | ERS784806 | 8791975 | 51 | 67.72 | 967784 | 557 | | | | |
| BCC1317 | 3 | ENVI , UK | ERS785017 | 8761762 | 63 | 67.93 | 552562 | 949 | | | | |
| BCC1622 | 3 | CF Isolate, USA | ERS784864 (GCA_900608515) | 8410623 | 51 | 68.05 | 615150 | 952 | | | | |
| BCC1623 | 3 | CF Isolate, USA | ERS784879 | 8430801 | 78 | 67.87 | 365793 | 953 | | | | |
| BCC1647 | 3 | *B. gladioli* pv. *gladioli*, LMG 6882, USA | ERS784943 | 8875626 | 51 | 67.7 | 561180 | 955 | | | | |
| BCC1649 | 3 | ENV; *B. gladioli* pv. *gladioli*, LMG 6880t4, Zimbabwe | ERS784812 | 8241275 | 51 | 68.3 | 554529 | 956 | | | | |
| BCC1651 | 3 | CF Isolate, LMG 18157 | ERS784844 | 7931757 | 44 | 68.29 | 595470 | 958 | | | | |
| BCC1668 | 3 | CF Isolate, UK | ERS784974 | 8456480 | 50 | 67.97 | 603154 | 966 | | | | |
| BCC1669 | 3 | CF Isolate, UK | ERS784982 | 8173618 | 52 | 68.14 | 485051 | 967 | | | | |
| BCC1677 | 3 | CF Isolate, USA | ERS785071 | 8109976 | 41 | 68.25 | 632433 | 972 | | | | |
| BCC1683 | 3 | CF Isolate, USA | ERS784962 | 8306965 | 44 | 68.17 | 886986 | 977 | | | | |
| BCC1687 | 3 | CF Isolate, USA | ERS784833 | 8185231 | 63 | 68.21 | 336464 | 981 | | | | |
| BCC1688 | 3 | CF Isolate, USA | ERS784848 | 8455786 | 75 | 68.14 | 319266 | 982 | | | | |
| BCC1691 | 3 | CF Isolate, USA | ERS784898 | 8419445 | 58 | 68.04 | 1964774 | 952 | | | | |

| | | | | | | | | | | | | |
|---|---|---|---|---|---|---|---|---|---|---|---|---|
| BCC1694 | 3 | CF Isolate, USA | ERS784946 | 8468446 | 45 | 67.9 | 688705 | 986 | | | | |
| BCC1696 | 3 | CF Isolate, USA | ERS784976 | 8266956 | 75 | 68.19 | 429819 | 988 | | | | |
| BCC1698 | 3 | CF Isolate, USA | ERS784818 | 8489077 | 68 | 68.1 | 434629 | 990 | | | | |
| BCC1700 | 3 | CF Isolate, USA | ERS784851 | 8488515 | 43 | 68.08 | 1416119 | 992 | | | | |
| BCC1703 | 3 | CF Isolate, USA | ERS785056 | 8286893 | 43 | 68.22 | 599795 | 955 | | | | |
| BCC1707 | 3 | CF Isolate, USA | ERS784948 | 8009488 | 42 | 68.22 | 534508 | 997 | | | | |
| BCC1709 | 3 | CF Isolate, USA | ERS785035 | 8457160 | 80 | 68.12 | 414686 | 998 | | | | |
| BCC1713 | 3 | CF Isolate, USA | ERS785074 | 8456910 | 192 | 68.05 | 147146 | 1001 | | | | BCC1713 |
| BCC1714 | 3 | CF Isolate, USA | ERS785080 | 8498126 | 70 | 68.08 | 361363 | 1002 | | | | |
| BCC1720 | 3 | CF Isolate, USA | ERS784985 | 7902991 | 43 | 68.29 | 770588 | 958 | | | | |
| BCC1721 | 3 | CF Isolate, USA | ERS784820 | 8216814 | 30 | 68.23 | 1289329 | 1006 | | | | |
| BCC1722 | 3 | CF Isolate, USA | ERS784836 | 7994327 | 63 | 68.18 | 346974 | 1007 | | | | |
| BCC1723 | 3 | CF Isolate, USA | ERS784853 | 8184213 | 44 | 68.25 | 1570053 | 1008 | | | | |
| BCC1724 | 3 | CF Isolate, USA | ERS785060 | 8510702 | 64 | 68.03 | 505438 | 1006 | | | | |
| BCC1725 | 3 | CF Isolate, USA | ERS784869 | 8559474 | 62 | 68.03 | 521790 | 1010 | | | | |
| BCC1726 | 3 | CF Isolate, USA | ERS784885 | 8129584 | 39 | 68.29 | 848284 | 1011 | | | | |
| BCC1727 | 3 | CF Isolate, USA | ERS785068 | 8153985 | 43 | 68.29 | 921754 | 1012 | | | | |
| BCC1728 | 3 | CF Isolate, USA | ERS785003 | 8123767 | 50 | 68.29 | 391569 | 1013 | | | | |
| BCC1729 | 3 | CF Isolate, USA | ERS784902 | 8365875 | 53 | 67.89 | 662719 | 1014 | | | | |
| BCC1730 | 3 | CF Isolate, Canada | ERS784917 | 8528922 | 44 | 68.12 | 793173 | 1015 | | | | |
| BCC1732 | 3 | CF Isolate, USA | ERS784933 | 8810912 | 64 | 67.67 | 606420 | 1017 | | | | |
| BCC1733 | 3 | CF Isolate, USA | ERS784951 | 8462420 | 53 | 68.03 | 519208 | 1018 | | | | |
| BCC1734 | 3 | CF Isolate, USA | ERS785079 | 8187286 | 59 | 68.24 | 757647 | 901 | | | | |
| BCC1754 | 3 | CF Isolate, USA | ERS1371583 | 8372702 | 104 | 68.2 | 416499 | Novel | | | | |
| BCC1755 | 3 | CF Isolate, USA | ERS1328941 | 8248941 | 166 | 68.2 | 324811 | Novel | | | | |
| BCC1757 | 3 | CF Isolate, USA | ERS1371584 | 8106595 | 108 | 68.25 | 281288 | Novel | | | | |
| BCC1758 | 3 | CF Isolate, USA | ERS1371585 | 8411592 | 135 | 68.16 | 347044 | Novel | | | | |
| BCC1760 | 3 | CF Isolate, USA | ERS1371586 | 8142112 | 101 | 68.21 | 351231 | 972 | | | | |
| BCC1763 | 3 | CF Isolate, USA | ERS1328942 | 8711025 | 150 | 68.08 | 409905 | Novel | | | | |
| BCC1765 | 3 | CF Isolate, USA | ERS1328883 | 8049558 | 131 | 68.1 | 267531 | Novel | | | | |
| BCC1767 | 3 | CF Isolate, USA | ERS1371590 | 8144838 | 89 | 68.28 | 365649 | Novel | | | | |
| BCC1772 | 3 | CF Isolate, USA | ERS1328773 | 8423448 | 169 | 68.03 | 340707 | Novel | | | | |

| | | | | | | | | | | | | |
|---|---|---|---|---|---|---|---|---|---|---|---|---|
| BCC1775 | 3 | CF Isolate, USA | ERS1333277 | 8368392 | 155 | 68.13 | 332589 | 992 | | | | |
| BCC1777 | 3 | CF Isolate, USA | ERS1328886 | 7915650 | 135 | 68.09 | 275756 | Novel | | | | |
| BCC1778 | 3 | CF Isolate, USA | ERS1328776 | 8287507 | 135 | 68.04 | 284651 | Novel | | | | |
| BCC1784 | 3 | CF Isolate, USA | ERS1328888 | 8106978 | 167 | 68.17 | 273802 | Novel | | | | |
| BCC1786 | 3 | CF Isolate, USA | ERS1328945 | 8316388 | 145 | 68.18 | 347752 | Novel | | | | |
| BCC1787 | 3 | CF Isolate, USA | ERS1371593 | 8239682 | 112 | 68.27 | 371296 | Novel | | | | |
| BCC1788 | 3 | CF Isolate, USA | ERS1371594 | 8626684 | 376 | 67.84 | 291652 | Novel | | | | |
| BCC1789 | 3 | CF Isolate, USA | ERS1328781 | 8320110 | 145 | 68.05 | 274262 | 901 | | | | |
| BCC1791 | 3 | CF Isolate, USA | ERS1328782 | 8138900 | 125 | 68.14 | 334966 | Novel | | | | |
| BCC1793 | 3 | CF Isolate, USA | ERS1328783 | 8137484 | 108 | 68.24 | 280591 | Novel | | | | |
| BCC1794 | 3 | CF Isolate, USA | ERS1328946 | 8325891 | 167 | 68.23 | 281967 | Novel | | | | |
| BCC1796 | 3 | CF Isolate, USA | ERS1328891 | 8374863 | 109 | 68.08 | 351151 | 902 | | | | |
| BCC1799 | 3 | CF Isolate, USA | ERS1328786 | 8102290 | 84 | 68.22 | 343906 | 1015 | | | | |
| BCC1800 | 3 | CF Isolate, USA | ERS1328892 | 7908681 | 135 | 68.29 | 257500 | Novel | | | | BCC1800 |
| BCC1801 | 3 | CF Isolate, USA | ERS1328787 | 7951550 | 97 | 68.24 | 316850 | 958 | | | | |
| BCC1802 | 3 | CF Isolate, USA | ERS1328947 | 8450656 | 130 | 68.11 | 270804 | Novel | | | | |
| BCC1806 | 3 | CF Isolate, USA | ERS1333279 | 8476169 | 174 | 68.05 | 273269 | 992 | | | | |
| BCC1807 | 3 | CF Isolate, USA | ERS1371599 | 8465961 | 109 | 68 | 304296 | Novel | | | | |
| BCC1808 | 3 | CF Isolate, USA | ERS1371600 | 7962624 | 122 | 68.22 | 293939 | 901 | | | | |
| BCC1809 | 3 | CF Isolate, USA | ERS1328791 | 8633417 | 162 | 67.87 | 235923 | Novel | | | | |
| BCC1810 | 3 | CF Isolate, USA | ERS1328948 | 8276123 | 94 | 68.22 | 310714 | 955 | | | | |
| BCC1811 | 3 | CF Isolate, USA | ERS1328792 | 8596479 | 208 | 67.93 | 305514 | Novel | | | | |
| BCC1813 | 3 | CF Isolate, USA | ERS1328793 | 8559601 | 130 | 68.01 | 293202 | Novel | | | | |
| BCC1816 | 3 | CF Isolate, Canada | ERS1328896 | 8179064 | 97 | 68.12 | 326051 | Novel | | | | |
| BCC1817 | 3 | CF Isolate, USA | ERS1328795 | 8110913 | 127 | 68.21 | 242936 | Novel | | | | |
| BCC1818 | 3 | CF Isolate, USA | ERS1333280 | 8154489 | 96 | 68.23 | 345879 | 1010 | | | | |
| BCC1822 | 3 | CF Isolate, USA | ERS1336063 | 8078754 | 104 | 68.18 | 402601 | Novel | | | | |
| BCC1823 | 3 | CF Isolate, USA | ERS1328798 | 8207755 | 107 | 68.31 | 336871 | Novel | | | | |
| BCC1826 | 3 | CF Isolate, USA | ERS1328949 | 8374984 | 115 | 68.13 | 338267 | Novel | | | | |
| BCC1827 | 3 | CF Isolate, USA | ERS1328800 | 8704353 | 169 | 67.88 | 269894 | Novel | | | | |
| BCC1830 | 3 | CF Isolate, USA | ERS1371602 | 8388969 | 115 | 68.05 | 419764 | Novel | | | | |
| BCC1833 | 3 | CF Isolate, USA | ERS1328803 | 8478257 | 126 | 68.02 | 390244 | Novel | | | | |

| | | | | | | | | | | | | |
|---|---|---|---|---|---|---|---|---|---|---|---|---|
| BCC1836 | 3 | CF Isolate, USA | ERS1328901 | 8169254 | 114 | 68.3 | 407323 | Novel | | | | |
| BCC1841 | 3 | CF Isolate, USA | ERS1328807 | 8713715 | 196 | 67.94 | 274939 | 955 | | | | |
| BCC1842 | 3 | CF Isolate, USA | ERS1328951 | 8157581 | 116 | 68.27 | 344808 | Novel | | | | |
| BCC1845 | 3 | CF Isolate, USA | ERS1328809 | 8127806 | 86 | 68.24 | 362515 | 902 | | | | |
| BCC1846 | 3 | CF Isolate, USA | ERS1371607 | 8405902 | 113 | 68 | 320953 | Novel | | | | |
| BCC1847 | 3 | CF Isolate, USA | ERS1371608 | 8398980 | 143 | 68.17 | 287220 | Novel | | | | |
| BCC1849 | 3 | CF Isolate, USA | ERS1371609 | 8211748 | 118 | 68.29 | 368142 | Novel | | | | |
| BCC1850 | 3 | CF Isolate, USA | ERS1371610 | 8519442 | 175 | 68.02 | 275270 | 977 | | | | |
| BCC1851 | 3 | CF Isolate, USA | ERS1371611 | 8365451 | 131 | 68.2 | 304291 | Novel | | | | |
| BCC1852 | 3 | CF Isolate, USA | ERS1371612 | 8412916 | 113 | 68.05 | 333375 | 902 | | | | |
| BCC1853 | 3 | CF Isolate, USA | ERS1328813 | 8212875 | 114 | 68.2 | 326166 | Novel | | | | |
| BCC1854 | 3 | CF Isolate, USA | ERS1371613 | 8895140 | 180 | 67.81 | 274690 | Novel | | | | |
| BCC1856 | 3 | CF Isolate, USA | ERS1328906 | 8038261 | 129 | 68.29 | 286277 | Novel | | | | |
| BCC1857 | 3 | CF Isolate, USA | ERS1371615 | 8229080 | 111 | 68.25 | 280379 | Novel | | | | |
| BCC1858 | 3 | CF Isolate, USA | ERS1371616 | 8186921 | 115 | 68.22 | 307394 | Novel | | | | |
| BCC1859 | 3 | CF Isolate, USA | ERS1371617 | 8251063 | 116 | 68.2 | 334836 | 1010 | | | | |
| BCC1860 | 3 | CF Isolate, USA | ERS1371618 | 8617021 | 136 | 68 | 347760 | 952 | | | BCC1860 | BCC1860 |
| BCC1862 | 3 | CF Isolate, USA | ERS1371620 | 8378040 | 108 | 68.14 | 294407 | Novel | | | | |
| BCC1863 | 3 | CF Isolate, USA | ERS1371621 | 8264387 | 121 | 68.23 | 294023 | Novel | | | | |
| BCC1866 | 3 | CF Isolate, USA | ERS1371622 | 8450941 | 143 | 67.99 | 287257 | 1002 | | | | |
| BCC1868 | 3 | CF Isolate, USA | ERS1371624 | 7929417 | 92 | 68.19 | 393583 | Novel | | | | |
| BCC1869 | 3 | CF Isolate, USA | ERS1328821 | 8594173 | 159 | 67.92 | 311968 | 902 | | | | |
| BCC1872 | 3 | CF Isolate, USA | ERS1371627 | 8415473 | 162 | 68.15 | 286407 | Novel | | | | |
| BCC1881 | 3 | ENV; *B. gladioli* pv. *agaricicola*, NCPPB 3580, UK | ERS1371629 | 8378078 | 125 | 68.2 | 350910 | Novel | | | | |
| BCC1882 | 3 | ENV; *B. gladioli* pv. *agaricicola*, NCPPB 3632, UK | ERS1371630 | 8374159 | 130 | 68.2 | 350910 | Novel | | | | |
| BCC1883 | 3 | ENV; *B. gladioli* pv. *agaricicola*, NCPPB 3852, New Zeland | ERS1371631 | 8116617 | 99 | 68.31 | 603194 | Novel | | BCC1883 | BCC1883 | BCC1883 |

**Table S2**. Primers and *B. gladioli* isolates used for generation of biosynthetic gene cluster insertional mutants

| **Target Cluster** | **Target gene** | **Strain** | **Forward Primer (5'-3')**[a] | **Reverse Primer (5'-3')**[a] | **Reference** |
|---|---|---|---|---|---|
| Gladiolin | *gbnD1* | BCC0238 | ATTTCTAGAGTTGCTGATC GACGCCTATC | TATGAATTCCGCCATAACC GAACGAACTC | Song *et al.* 2017 |
| Toxoflavin | *toxA* | BCC0238 | GACATCTAGATTGGTGATG TTTCCGGCAAG | GGCGGGTACCGAACACGT CCCAGAAACCAG | This work |
| Bongkrekic acid | *bonA* | BCC1710 | ATTTCTAGAAGTATCCGCA TTTTCGTCGC | TATGAATTCGATCGATCAG TTGCGCTTCC | This work |
| Caryoynencin | *cayA* | BCC1697 | GCTCTAGACATGTCGTGAT AAGGCGGC | GCGGAATTCACCGTGAGC TATGCCGAG | This work |

**Footnote**: [a] The underlined primer sequence indicates the restriction enzyme site overlap used for cloning the PCR products into pGpΩTp

**SUPPLEMENTARY FIGURES**

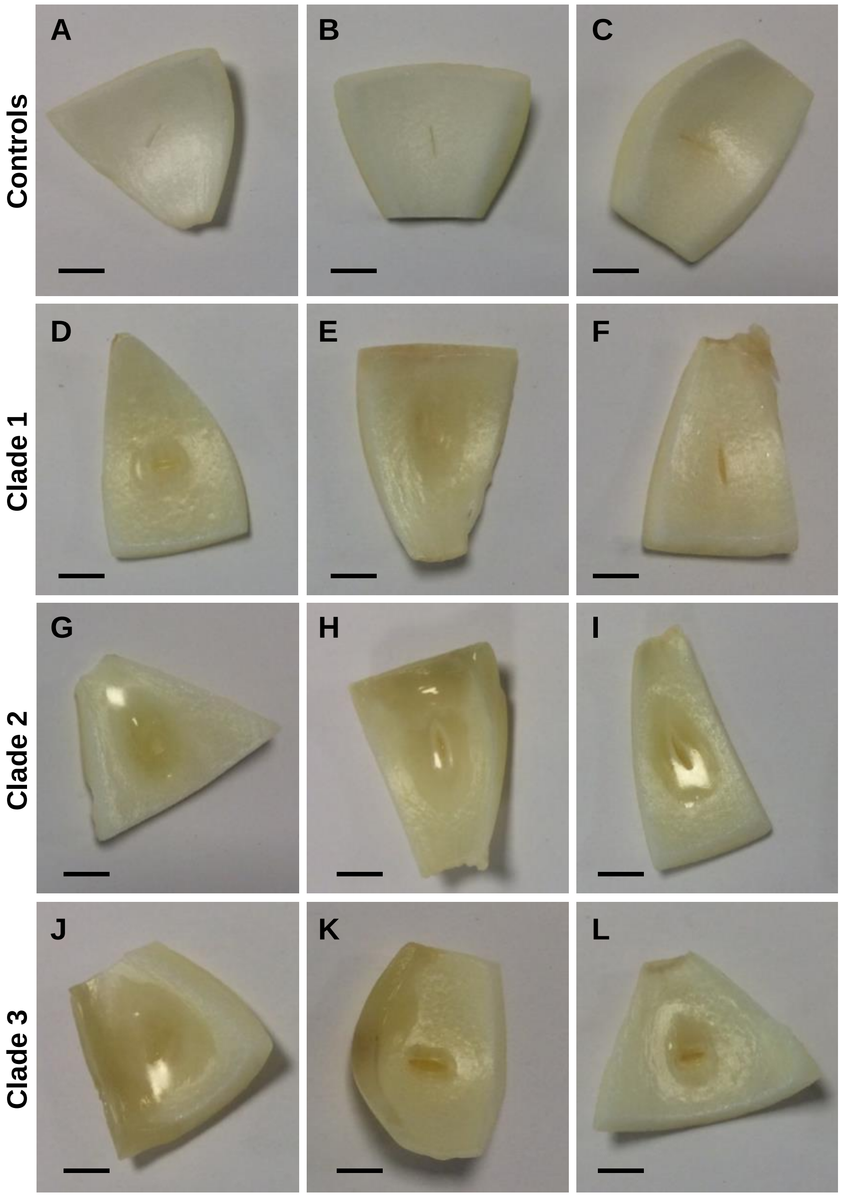

**Figure S1. *B. gladioli* from all evolutionary clades are capable of onion rot.** The ability of *B. gladioli* strains to degrade onion tissue was tested by inoculating injured onion tissue with standardized bacterial cultures. Inoculated mushrooms were incubated at 30°C for 48 hours with the following shown in each panel (each row are either controls or *B. gladioli* clades as indicated on the left): (A) No treatment control, (B) TSB only, (C) *E. coli* NCTC 12241, (D) *B. gladioli* Clade 1A strain BCC1710, (E) Clade 1B strain BCC1675, (F) Clade 1C strain BCC1678, (G) *B. gladioli* Clade 2 strain BCC1731, (H) *B. gladioli pv. allicola* reference Clade 2 strain BCC1645, (I) *B. gladioli pv. allicola* reference Clade 2 strain BCC1646, (J) *B. gladioli* Clade 3 strain BCC0238, (K) *B. gladioli pv. agaricola* reference Clade 3 strain BCC1883 (NCPPB 3852), (L) *B. gladioli pv. gladioli* reference Clade 3 strain BCC771 (LMG 2216$^{T}$). Pitting and tissue degradation was apparent in all *B. gladioli* inoculated mushrooms; a scale bar (1 cm) is shown in each panel to enable comparison.

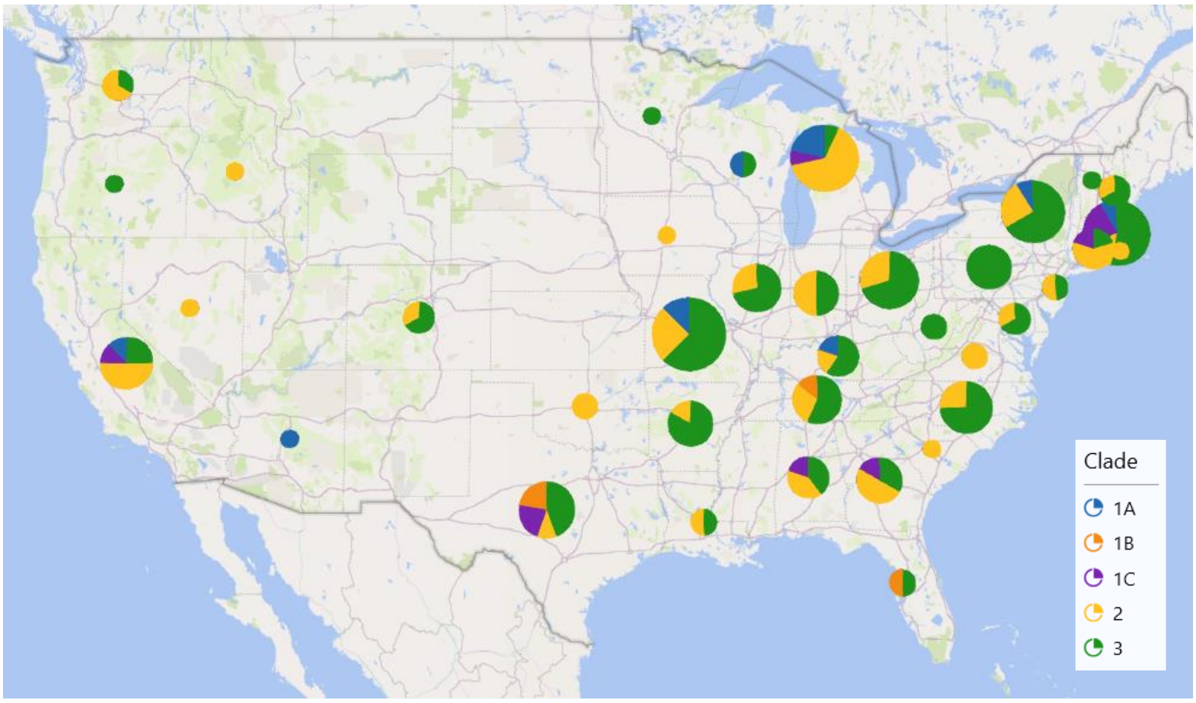

**Figure S2. CF infection associated with each *B. gladioli* clade is widely distributed across the United States of America.** The state location of the submitting CF treatment centre for 181 CF *B. gladioli* strains in the collection are shown. The *B. gladioli* clade type was plotted against geographic location using the Power Map tool in Microsoft Excel Office 365. The circle size is proportional to the total number of strains at each location and coloured segments indicate the proportion of each clade present (see key).

**Figure S3. UHPLC-ESI-Q-TOF-MS analysis of metabolites produced by *B. gladioli* isolates**. Structures, calculated masses, extracted ion chromatograms (EICs) and high resolution mass spectra are displayed for each of the following compounds (corresponding strain): (a) toxoflavin (BCC1883); (b) enacyloxin IIa (BCC1883); (c) bongkrekic acid (BCC1710); (d) icosalide A1 (BCC0238); (e) gladiolin (BCC0238); (f) sinapigladioside (BCC1883); and (g) caryoynencin (BCC1883). The MS spectra correspond to the region underneath the peak with an asterisk in each instance.

### (a) Toxoflavin

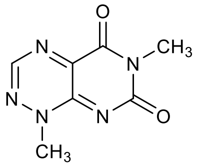

**Toxoflavin**
Chemical Formula: $C_7H_7N_5O_2$
Exact Mass: 193.0600
$[M+H]^+$: 194.0673
$[M+Na]^+$: 216.0492

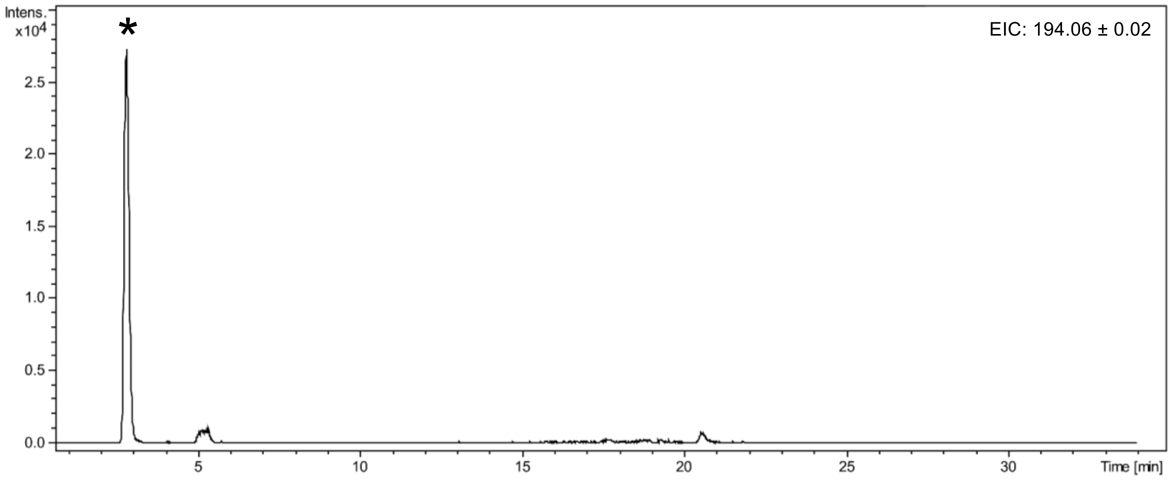

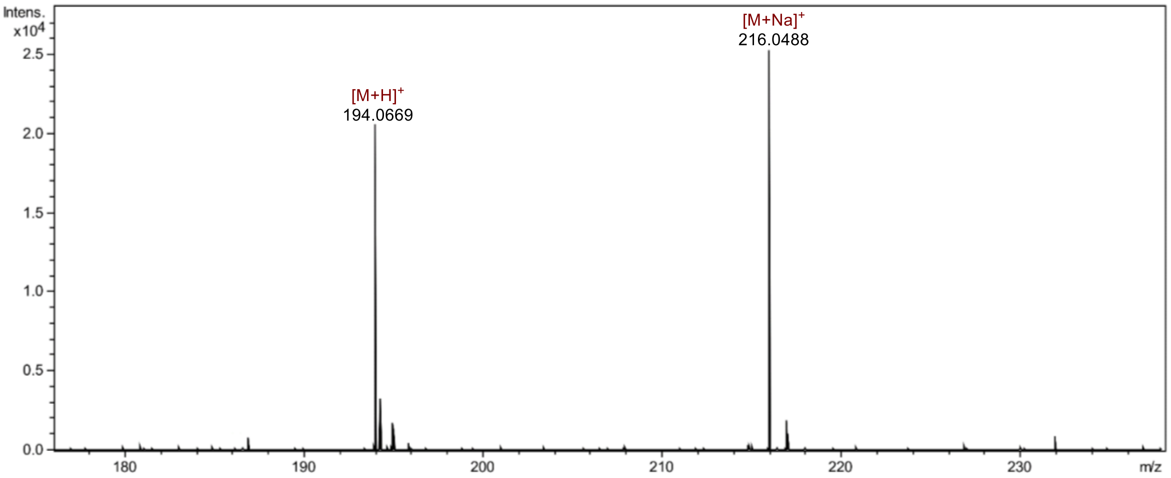

### (b) Enacyloxin IIa

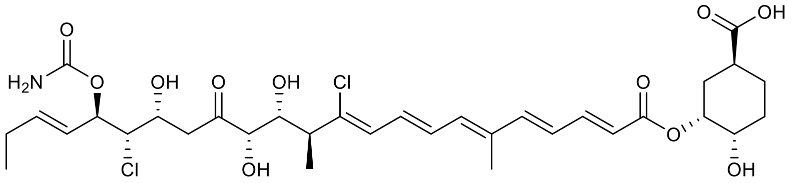

**Enacyloxin IIa**
Chemical Formula: $C_{33}H_{45}Cl_2NO_{11}$
Exact Mass: 701.2370
$[M+H]^+$: 702.2442
$[M+Na]^+$: 724.2262

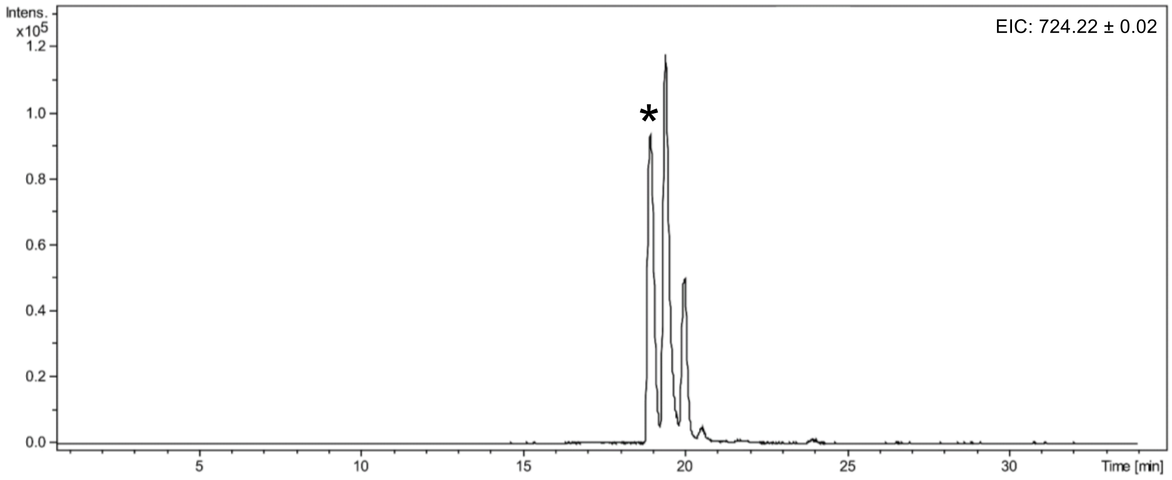

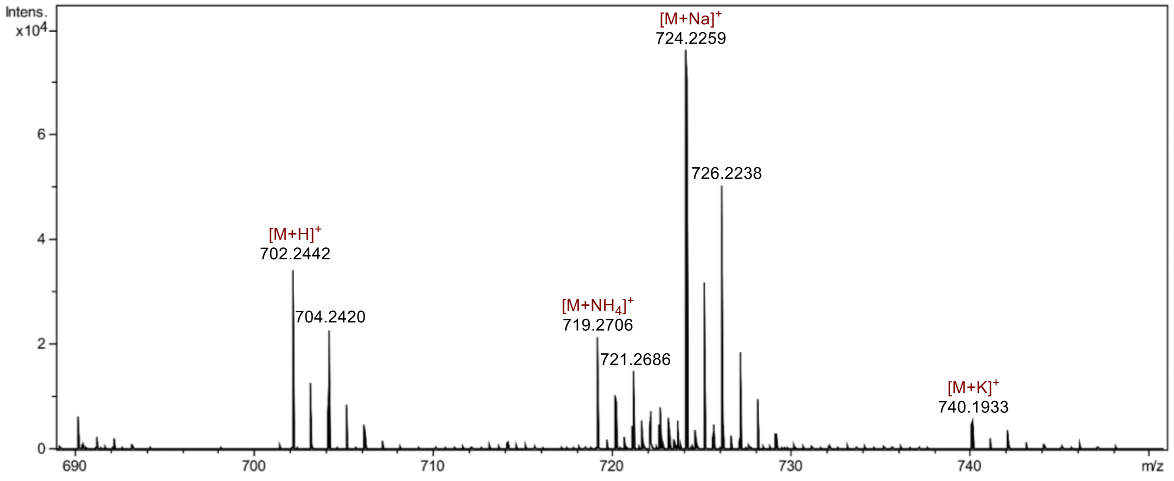

### (c) Bongkrekic acid

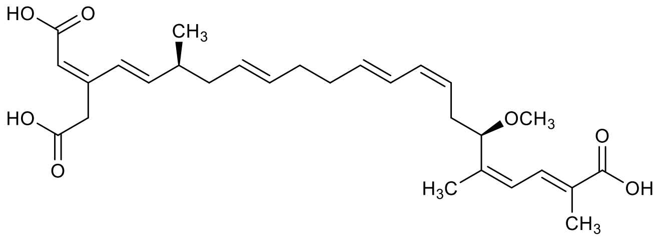

**Bongkrekic Acid**
Chemical Formula: $C_{28}H_{38}O_7$
Exact Mass: 486.2618
$[M+H]^+$: 487.2690
$[M+Na]^+$: 509.2510

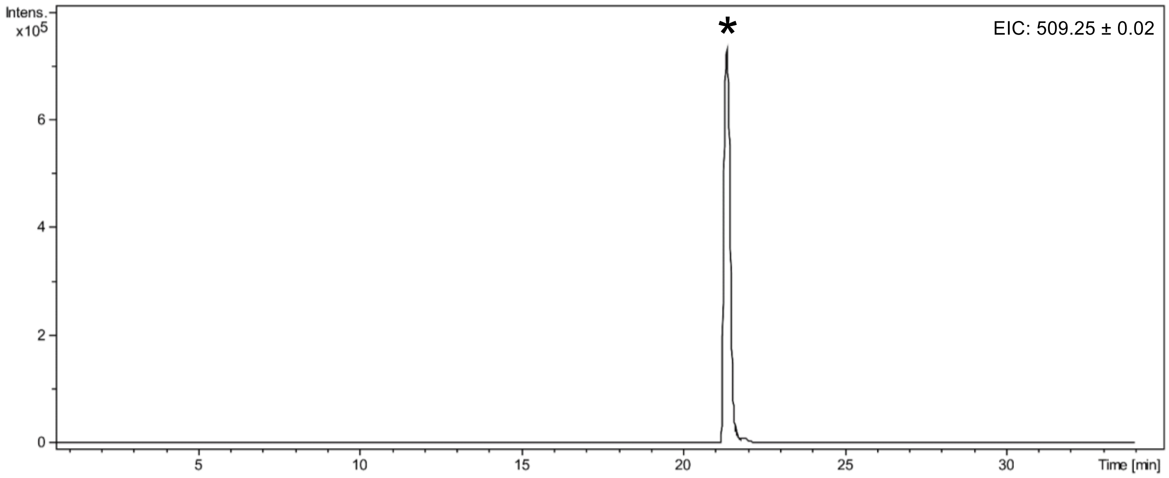

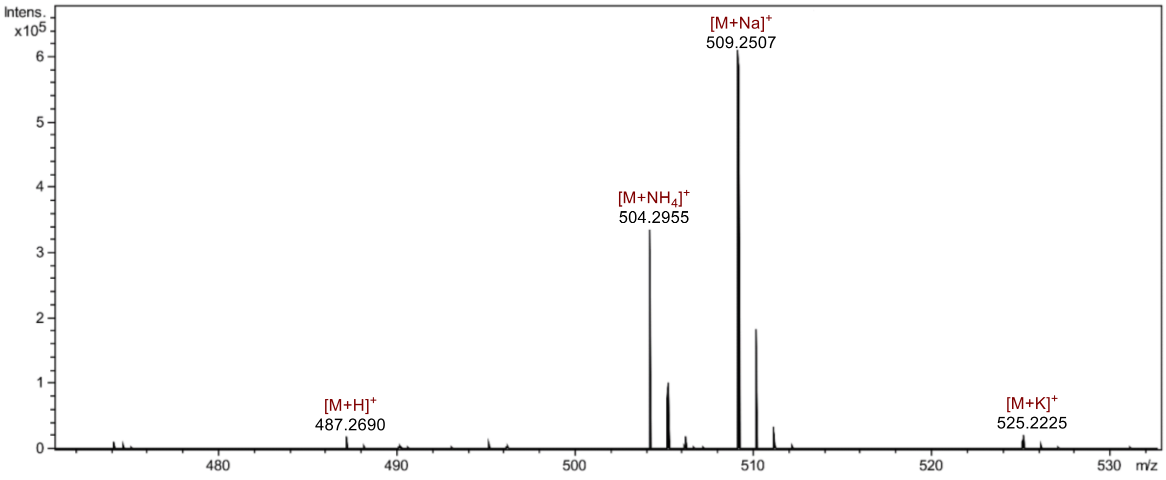

### (d) Icosalide A1

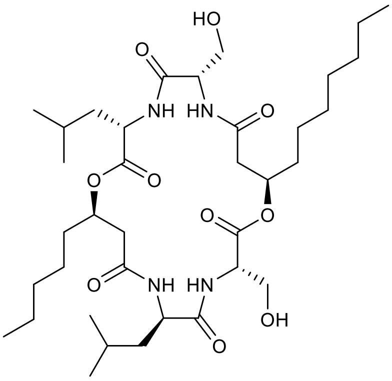

**Icosalide A1**
Chemical Formula: $C_{36}H_{64}N_4O_{10}$
Exact Mass: 712.4622
$[M+H]^+$: 713.4695
$[M+Na]^+$: 735.4515

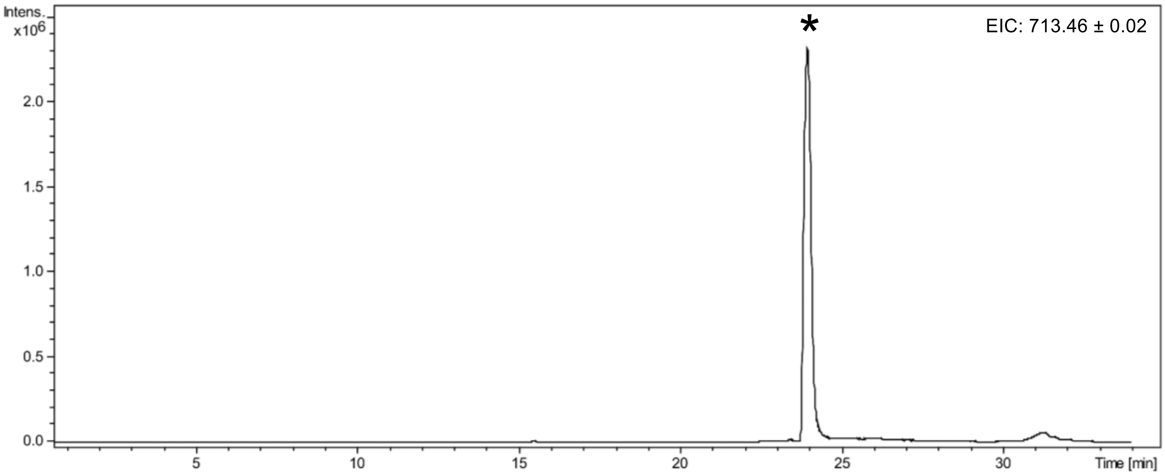

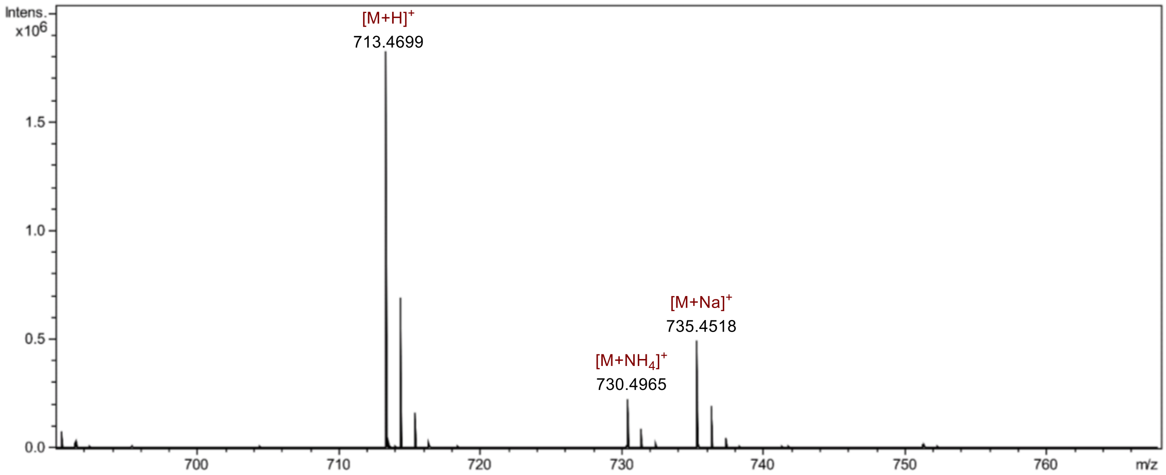

### (e) Gladiolin

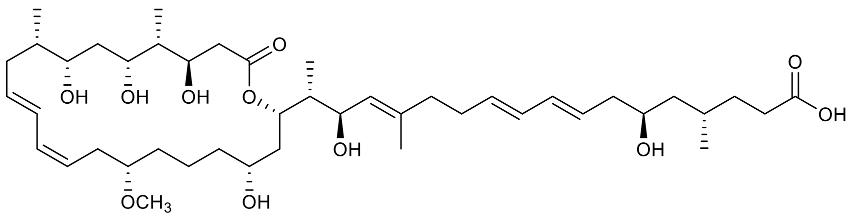

**Gladiolin**
Chemical Formula: $C_{44}H_{74}O_{11}$
Exact Mass: 778.5231
$[M+H]^+$: 779.5304
$[M+Na]^+$: 801.5123

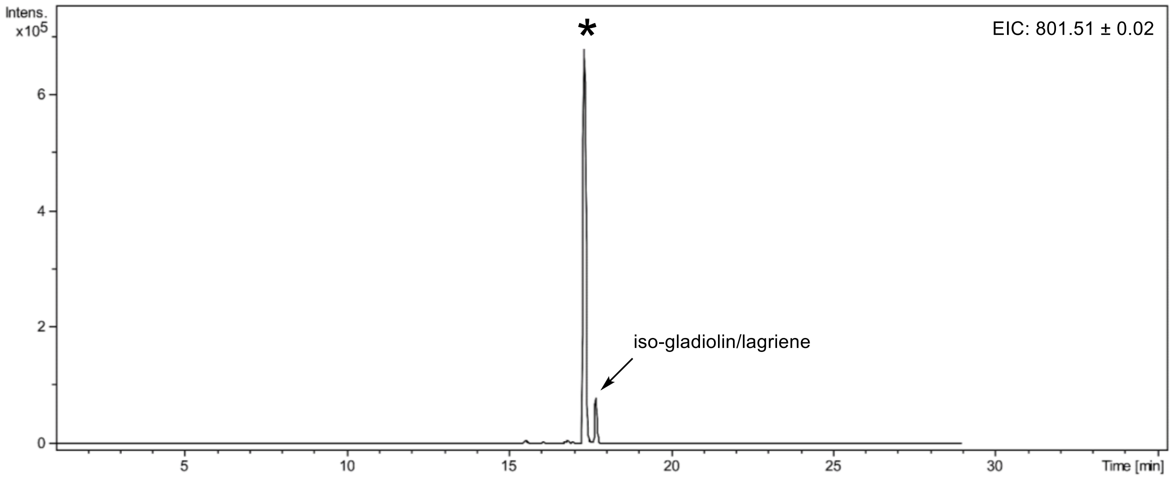

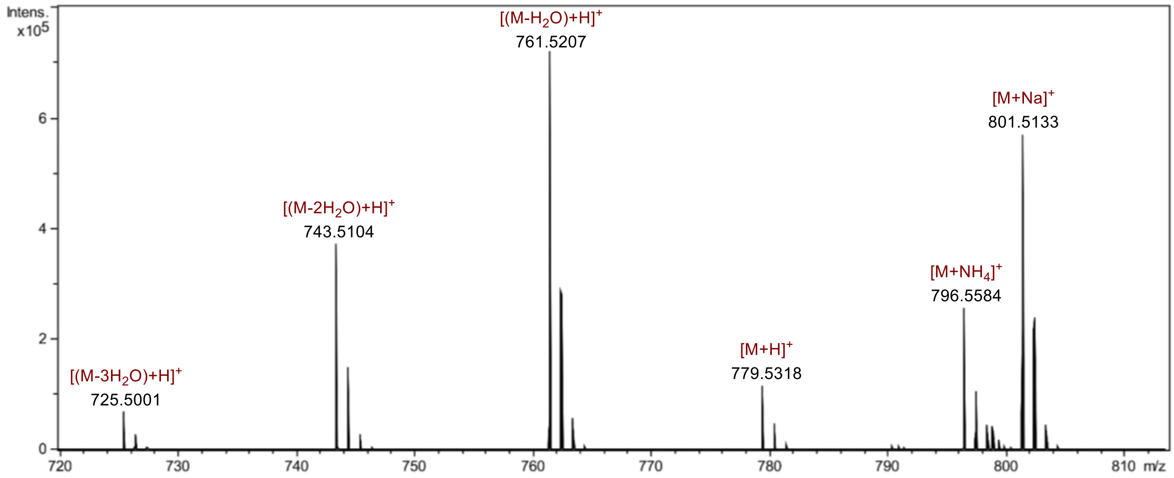

### (f) Sinapigladioside

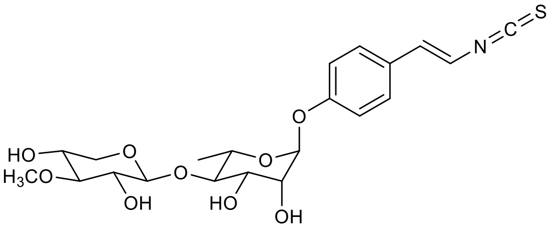

**Sinapigladioside**
Chemical Formula: $C_{21}H_{27}NO_9S$
Exact Mass: 469.1407
$[M-H]^-$: 468.1334
$[(M+Na)-2H]^-$: 490.1153

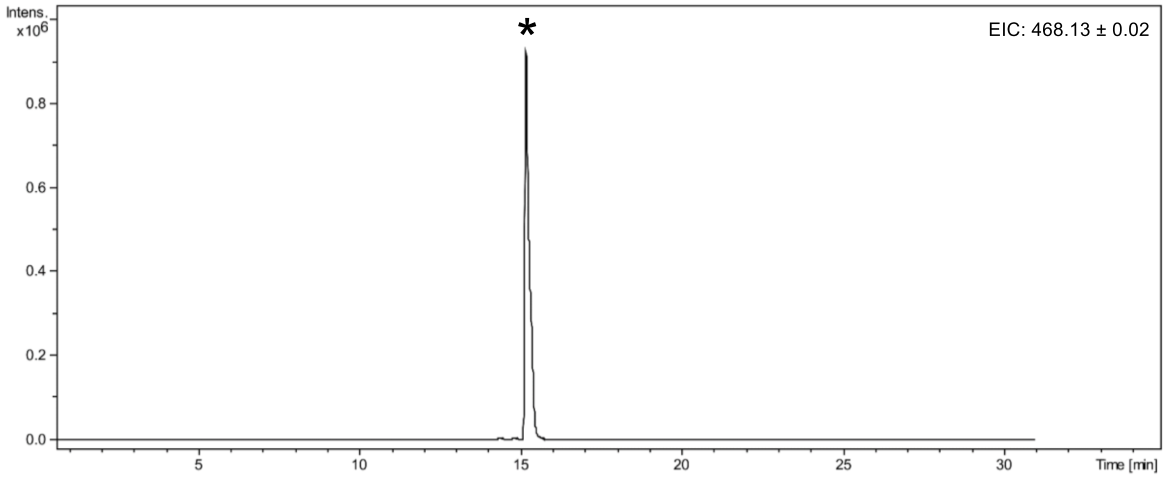

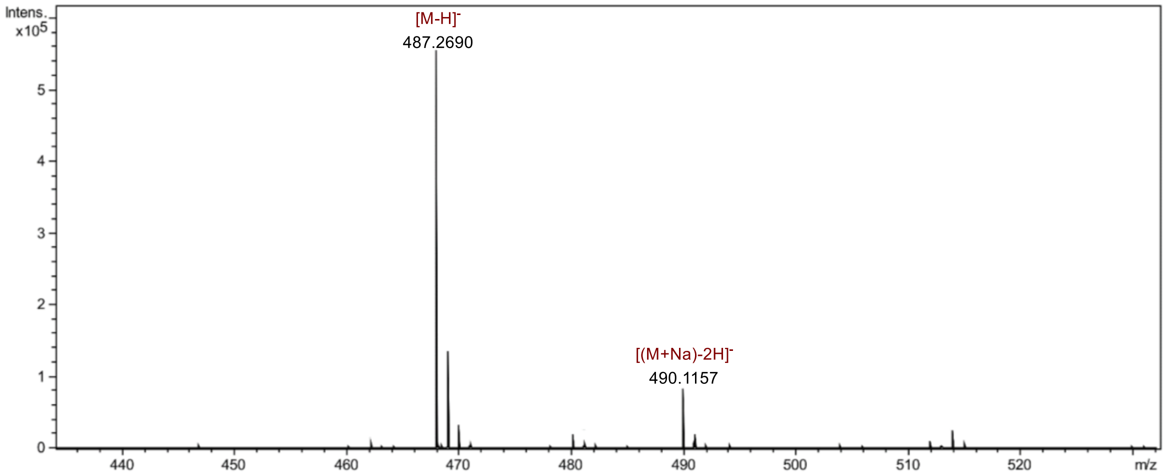

### (g) Caryoynencin

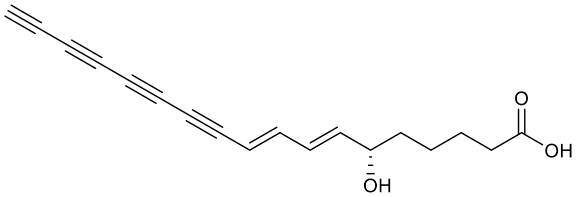

**Caryoynencin**
Chemical Formula: $C_{18}H_{16}O_3$
Exact Mass: 280.1099
$[M-H]^-$: 279.1027
$[(M+Na)-2H]^-$: 301.0846

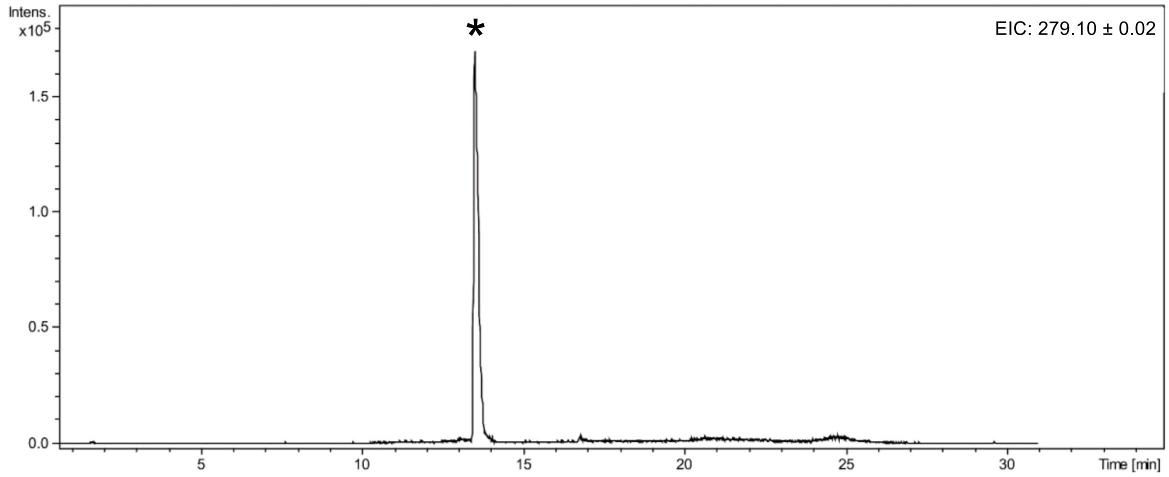

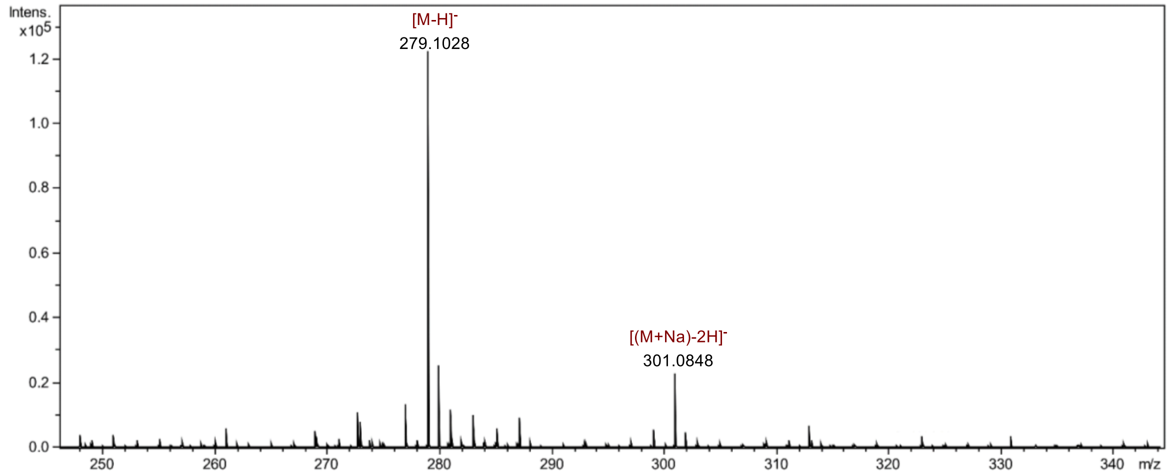

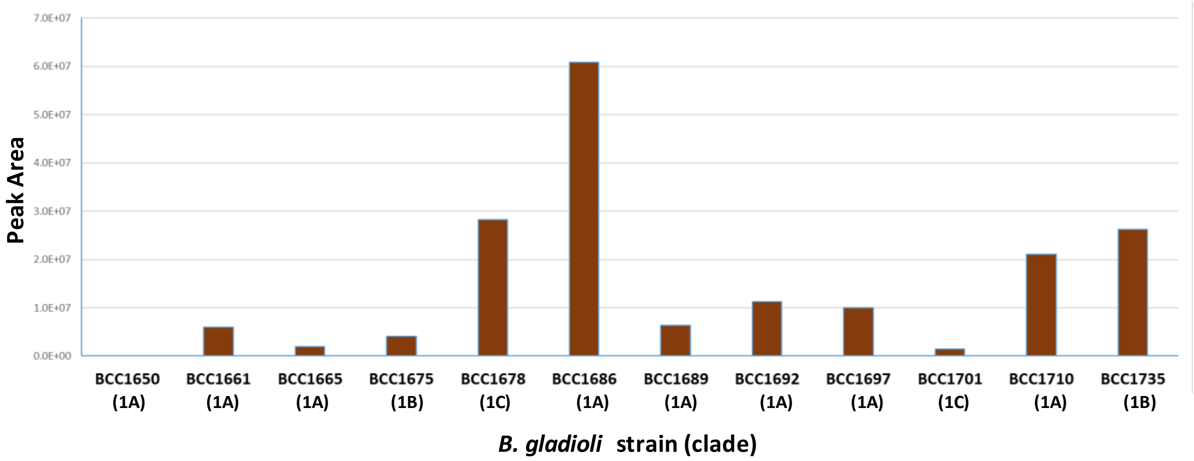

**Figure S4. *B. gladioli* clade 1 strains have a variable ability to produce bongkrekic acid.** Production of the toxin was evaluated by quantitative HPLC analysis (the mean peak height is plotted) of spent agar extracts analysed after 3 days growth of *B. gladioli* at 37$^{o}$C on BSM-G medium. The mean peak height for triplicate extracts is plotted for: 8 clade 1A, 2 clade 1B, and 2 clade 1C strains (isolate names and clade are indicated). Peak height was measured at 210-400 nm.

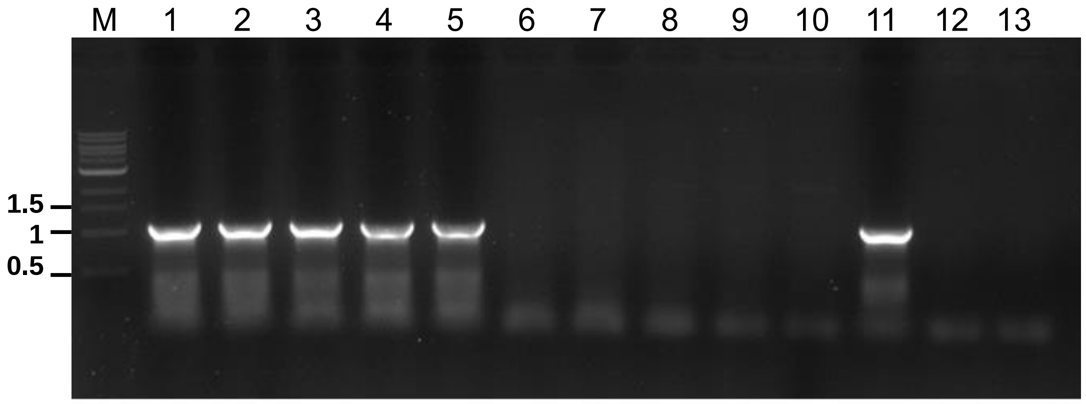

**Figure S5. Detection of *B. gladioli* clade 1 strains by bongkrekic acid *bonA* gene PCR.** An example PCR to detect the *bonA* gene of toxin positive clade 1 *B. gladioli* strains is shown with lanes as follows: M, molecular size markers (relevant size in kb are shown); 2 to 5 are bongkrekic acid positive strains BCC1710, 1650, 1661, 1735 and 1701, respectively; 6 to 10 are bongkrekic negative strains BCC1622, a colony variant 1622c2, 1723, 1620 and 1662, respectively; 11 is the positive pathovar control strain *B. gladioli* pv. *cocovenenans* LMG 18113; 12 is a negative control strains BCC0238 from clade 3; and lane 13 is the water PCR blank.

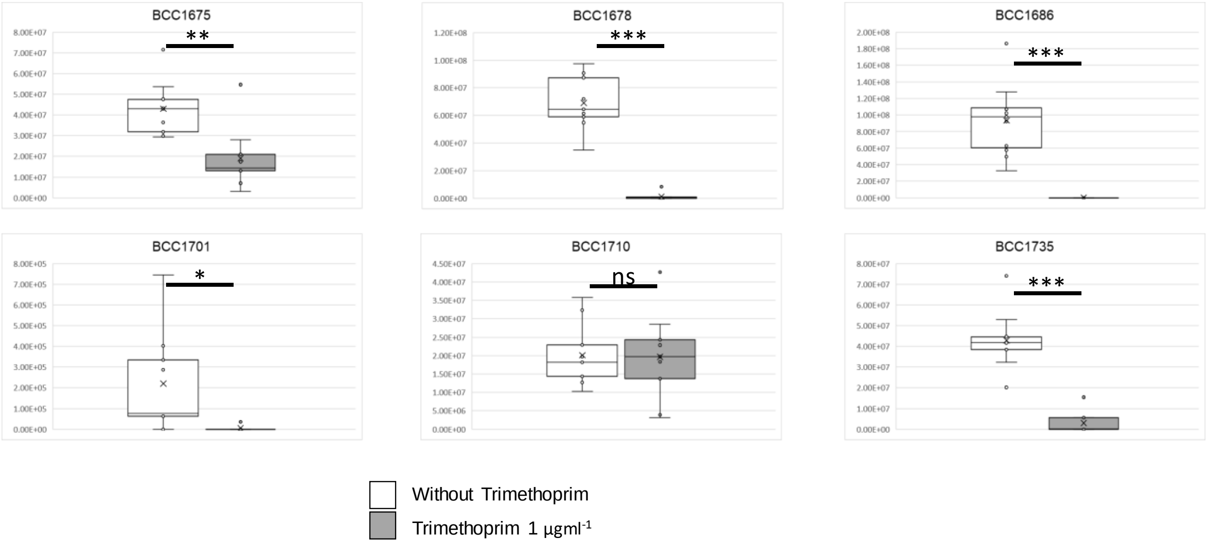

**Figure S6. Subinhibitory trimethoprim reduces bongrekic acid production.** HPLC analysis of bongkrekic acid production in 6 group 1 *B. gladioli* is shown (clade 1A, BCC1686 and BCC1710; clade 1B, BCC1675 and BCC1735; and clade 1C, BCC1678 and BCC1701). Bongkrekic acid production in the absence (white boxes) and presence (1 µg/mL; grey boxes) of the antibiotic trimethoprim is shown (n=9). Significant suppression (*$p<0.05$); **$p<0.01$; and ***$p<0.001$) of toxin production in the presence of trimethoprim occurred in 5 strains, with non-significant (ns) change observed for strain BCC1710. Peak height measured at 210-400 nm.
